# Exosomes isolated from two different cell lines using three different isolation techniques show variation in physical and molecular characteristics

**DOI:** 10.1101/2020.06.06.122952

**Authors:** Manish Dash, Kanagaraj Palaniyandi, Satish Ramalingam, S. Sahabudeen, N S Raja

## Abstract

Exosomes are the nanoscopic lipid bi-layered extracellular vesicles with the potential to be utilized as targeted therapeutics. In our investigation, we compared three major exosome isolation techniques that were Total Exosome Isolation reagent (TEI), Protein organic solvent precipitation (PROSPR) and differential ultracentrifugation (UC) based on the biophysical and physicochemical characteristics of exosomes isolated from COLO 205 and MCF-7 cancer cell’s conditioned media with an aim to select a suitable method for translational studies. 3D image analysis and particle size distribution of exosomes from their HRTEM images depicted the morphological differences. Molecular and analytical characterization of exosomes using western blotting, Raman and ATR-FTIR spectroscopy and the multivariate analysis on the spectral data obtained, assessed for better molecular specifications and purity of particle. TEI method isolated exosomes with higher exosomal yield, purity, and recovery directly translatable into drug delivery and targeted therapeutics whereas ultracentrifuge had good recovery of particle morphology but showed particle aggregation and yielded exosomes with smaller mean size. PROSPR technique isolated a mixture of EVs, showed lower protein recovery in PAGE and western blotting but higher spectroscopic protein to lipid ratio and distinguishable EV population in multivariate analysis compared to exosomes isolated by TEI and UC. This comparative study should help in choosing a specific exosome isolation technique required for the objective of downstream applications.

## 1. Introduction

Exosomes are the minuscule ultrafine bioparticles, the lipid vesicles with a potential to carry cellular cargo from one cell to another^1,2^. They are involved in cell–cell communication playing a major role in growth and development, immune response and even in regulating the tumor microenvironment and disease progression^3–6^. Exosomes are efficient and robust in their function and have versatile cargo composed of uniquely sorted material carrying it from the cytosol of parent cell to the target cell throughout the body^7^. They are highly efficient due to their non – immunogenic delivery, their ability to easily fuse with the cell membrane, avoid phagocytosis and circumventing the lysosomes via bypassing the engulfment^8,9^.

Exosomes are the ideal drug delivery system with the applications of targeted therapeutics, prognosis and diagnostics of the diseases^9,10,11^. This is because the synthetic lipid nanoparticles like lipoplexes can only be used *in vitro* and not *in vivo* as a delivery system for gene therapy and transfection^12–14^. Also, in animal models, these lipoplexes elucidate a strong immune response with increased toxicity levels and accumulation in the liver, hindering the drug activity in the target^15,16^.

The efficiency, biocompatibility and low toxicity levels are been harnessed for multiple novel therapeutic strategies^17–21^. During the development of exosome-based therapeutics and diagnostics, it’s important to keep in mind about the biophysical and molecular characteristics of these particles^22–24^. The attributes of these particles like their morphology and topology, the size of the particle and its uniformity, optimization of the storage conditions and the mode of the delivery of particles into the target cells with improvement in the medicament potential of these particles play important role in engineering and targeting the exosomes for the treatment and therapy^25,26^. Isolation method of exosomes that can give a higher yield with better purity and recovery of the exosomes play a crucial role in the collection of high-quality exosomes for scaling up the operations in the industry^27–29^. There are several techniques already available and many more are in development. Techniques like Ultracentrifugation, density gradient purification, ultrafiltration, qEV isolation system, precipitating reagents, exosome capture by diamagnetic beads, lab on a chip and several magnetic nanoparticle-based techniques have been developed^25,30,31^. Size exclusion chromatographic technique involving specialized columns isolate exosomes with precision^32^. Patented and commercialized products like qEV (IZON), ExopureTM (Biovision), Exo - spinTM columns (CELL guidance systems) are been used in studies for the isolation. Immunoaffinity based techniques like ELISA, immunomagnetic nanotechnology-based approaches or diamagnetic beads all of them get coated with exosome-specific antibodies and conjugating chemistry for exosome capture^33–38^. Several exosomes precipitating reagents like like Total Exosome Isolation Reagents for serum, plasma and cell culture media (Invitrogen), ExoQuickTM kit and range of Exosome isolation reagents (SBI System Biosciences), 101 Bio (Fisher Scientific), MagCaptreTM Exosome Isolation Kit PS (FUJIFILM WAKO Pure Chemical Corporation), Exosome Purification and RNA Isolation Kits (Norgen Biotek Corp) etc have been developed for exosome isolation with quick and easy steps^29,39^. Lab on Chip techniques (LOC) have also come to picture for exosome isolation with ease and better bioseparation modules^40,41^. But some of these techniques are either very costly or yield less exosomes. SEC columns cost a lot, immunoaffinity and nanotechnology techniques take in low amount of sample and cost high. Other methods like LOC currently are still under process of development and optimization and only small amount of sample can be processed. In this case growing labs in developing countries need techniques that can suitably help them in exosome research. Need of techniques that can be cost effective while yielding good quality or exosomes in large number is crucial for translational research and clinical setup.

In our present study, we compared the differential physical and molecular characteristics of exosomes isolated using three different isolation protocols namely, the isolation via Total exosome isolation reagent from Invitrogen (TEI) (which had been proven to be the best exosome precipitating reagent among others present commercially by a previous study)^27^, via Protein Organic Solvent Precipitation method (PROSPR) (which is studied and applied very less in case of EV isolation) and via Differential Ultracentrifugation (UC) (the conventional technique). The exosomes were isolated from two metastatic cell lines, COLO 205 a colon adenocarcinoma and MCF-7 a human breast cancer cell line. The exosomes from both the cell lines were isolated from their serum-free conditioned media and were quantified for their yield and proteome. The exosomes were characterized by HRTEM and the images were analyzed for the recovery of morphology and distribution of particles, SDS PAGE analysis was done to observe the difference in their proteomic profiles and Western Blotting was done for molecular characterization using CD63 marker. Spectroscopic techniques were employed to study the exosomes for quick label free characterization and differentiation purpose in cost effective manner compared to downstream MS analysis and expression analysis. Raman Spectroscopy and ATR-FTIR were performed for fingerprinting of the molecular components of exosomes respectively. Statistical analysis and multivariate analysis of the Raman and ATR-FTIR spectra were done to check for the variation in components of spectra of the exosomes isolated from two different cell line sources using three different methods. Our investigation provided the basis to select the suitable protocol for isolating exosomes, having higher yield and better recovery of exosomes with ideal morphology that could be applied to drug delivery and targeted therapeutics.

## 2. Materials and Methods

### Cell culture and isolation of exosomes

COLO 205 colon adenocarcinoma cells and MCF-7 human breast cancer cells were culutured in high glucose DMEM (HiMedia^®^, India), supplemented with 10 % FBS (US Heat Inactivated HiMedia^®^, India) and 2% penicillin/streptomycin (antibiotic antimycotic solution HiMedia^®^, India). The cells were incubated at 37°C with 5% CO_2_ supply and were allowed to grow until 70% confluency. The cells were replaced in FBS depleted media and allowed to grow up to 85-90% confluency in FBS free media from which exosomes were isolated.

The isolation of the exosomes was performed via three different methods namely, Total Exosome Isolation reagent (TEI) (Invitrogen, USA), Protein Organic Solvent Precipitation i.e. acetone precipitation mediated (PROSPR) and Differential Ultracentrifugation (UC). The protocols used for the isolation by precipitation based TEI reagent was taken from the TEI Invitrogen manufacturer’s manual, the PROSPR method taken was developed by Xavier Gallart-palau et al^42^ and UC was taken from the manuscript of <u>Clotilde Théry</u> et al^43^.The schematics for step by step protocol is given in figure2.1 below.

**Figure 2.1.**
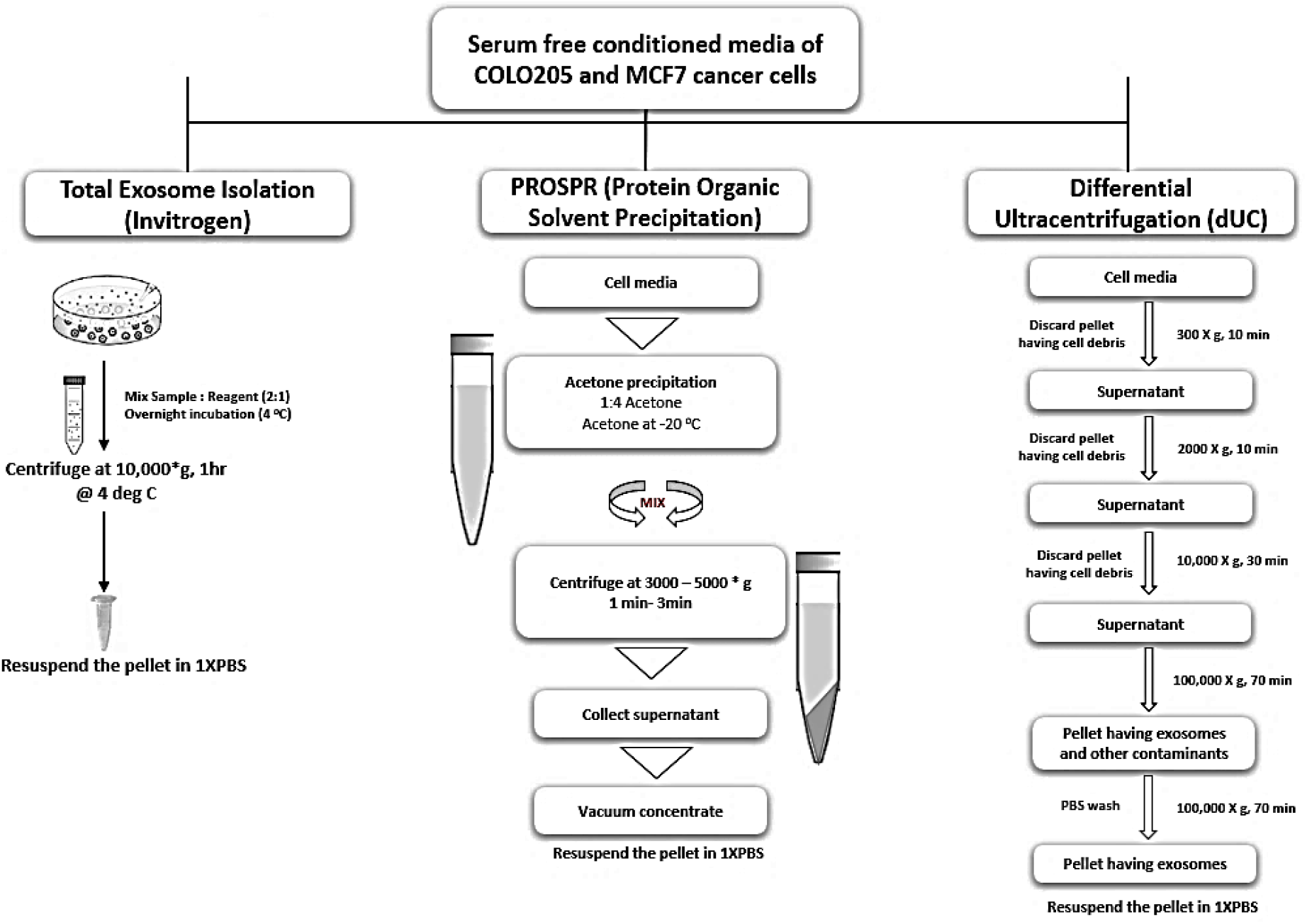
The three isolation techniques used in isolation of exosomes from serum-free conditioned media of COLO 205 and MCF-7 cancer cells were namely, TEI the method where the media to be processed was taken and mixed with the reagent in 2:1 ratio, vortexed properly and incubated overnight at 4°C. The exosomes were pelleted down at 10,000g for 60 min at 4°C according to manufacturer’s manual and were resuspended in 100μl of 1 X PBS for further studies, PROSPR the technique in which the conditioned media was mixed with pre-chilled ice-cold acetone in a 1:4 ratio and vortexed. This solution was centrifuged at 3000g for 2 min and the supernatant was collected and concentrated in a vacuum concentrator in vacuum-alcoholic (V-AL) mode (Eppendorf plus, Eppendorf, Hamburg, Germany). The concentrated crystals were resuspended in 100μl of 1 X PBS and Nuclease Free Water (NFW). The ultracentrifugation of the samples was carried out with a TLA-055 rotor of Beckman Coulter Optima MAX-XP ultracentrifuge (California, USA) by spinning them in lower centrifugal speed at 300g and gradually increasing the speed to 2000g and to 10,000g consecutively removing the cell debris in the form of a pellet. The supernatant collected from this stage was spun at 1,00,000g twice using ultracentrifuge at for 70 min each to isolate exosomes in the pellet at the final step. The first spin at ultra- high speed was done to remove the bigger vesicles. The supernatant was discarded and the pellet was washed with 1 X PBS in the second spin. The pellet (almost invisibly small in size) were resuspended in 100 μl of 1 X PBS and NFW.

One set of the samples of COLO 205 and MCF-7 conditioned media, were treated with 1X Proteinase K and incubated at 37°C for 30 min. The proteinase K activity was inactivated by incubation at 60°C for 10 min. This was done to check for the reduction of extracellular protein debris. The conditioned media were then processed further using the above three techniques for the isolation of exosomes.

### Quantification of exosome yield and exosome proteome

The quantification of the yield of exosomes and its total protein content was performed by Bradford’s reagent*, HiMedia® India.* The quantification of the total protein content was done by digesting the exosomes with RIPA buffer described by *Alcaraz C et al^44^.* The absorbance was read at 595nm in UV-Vis spectrophotometer (Jasco V-730 UV-Vis spectrophotometer, USA).

The quantification of Proteinase K treated exosome samples and their proteome content was done via the same procedure as above stated. All the experiments were done in triplicates and the results were analysed using OriginLab Pro 2020 ver. 9.4.

### High-Resolution Transmission Electron Microscopy

Jeol TEM – 2100 plus (Tokyo, Japan) was used for imaging of the exosomes. The imaging was done at high-resolution TEM with a magnification scale ranging from 20nm to 1μm with an applied voltage of 220 kV. For imaging, the samples were diluted 1:50 times and 2μl from the diluted working stock was made to set on a carbon-coated copper grid. No fixation or negative staining steps were performed for sample preparation. The grid was incubated overnight at 37°C with a desiccant kept in a closed chamber and was taken to the facility the next day for imaging.

### Image Analysis of the HRTEM images obtained for Exosomes

The 2D contour maps and 3D surface plots of exosome images obtained from HRTEM were used to create particle size distribution plots for understanding the morphology, size variation, and distribution of exosomes. The images were prepared in Fiji ImageJ software for the particle size distribution analysis by calibrating their scale, converting the images into an 8-bit grayscale type and adjusting the threshold values for binarization and particle tagging. 2D contour maps for calculation of particle parameters were generated in OriginLab software where the parameters were extracted from the contour line data matrix of the bitmap image created. The particles were then analysed for their area by calculating their Feret or Caliper diameter and Sauter mean diameter with normalized parameters and bin range. The 3D surface plots were modelled via Interactive 3D surface plots plugin and analysis manager in Fiji ImageJ software by optimizing the ratio of f(X, Y) to Z with 100% surface maxima and 0% surface minima. The size of the data grid was adjusted to the grid size of 512 nm sq. area with a perspective angle of 0.2° with optimum smoothing and z-scale enough to distinguish between individual exosomes and noise. The mode of simulation was set such that all the pixels of the image were connected without leaving holes. All the plots were computed by optimizing the parameters and plugin calling from the macro in consol. The graphing and analysis was done using OriginLab Pro 2020 ver. 9.4.9.

### SDS - PAGE and Western Blotting

Exosome samples were lysed with 1 X RIPA buffer at 70°C for 10 min followed by protein precipitation with ice-cold acetone in −20°C for 2 hours. The proteins were pelleted down in 0°C for 15 min 10,000 rpm. The pellets were air-dried and then digested with 10μl of 1 X Laemmli’s buffer for total protein denaturation and loading into PAGE. SDS PAGE under reducing condition was performed with 10 % separating gel having 1.5 M Tris pH 8.8 and 30% Acrylamide and 5 % stacking gel with 1 M Tris pH 6.8 and 30% Acrylamide at 100V. The gels were stained with Coomassie brilliant blue R-250 (Himedia^®^ India) for visualization of protein profile.

Molecular characterization of exosomes using their transmembrane protein marker CD63 was achieved via western blotting. Post electrophoresis, proteins were transferred to PVDF membrane in the Bio-Rad blot module with the help of transfer buffer. The antigens were probed with anti-CD63in 1:1000 dilution (Abcam, UK) primary antibody and rabbit anti-mouse secondary antibody in 1:2000 dilution (Abcam, UK) conjugated with HRP in phosphate buffered saline with added tween 20 (PBST) blocking buffer having 5% non-fat milk. Equal volumes of peroxide solution and luminol enhancer solution (Bio-Rad Clarity™ western ECL substrates, USA) were added before the visualization in ChemiDoc XRS+ gel documentation system (Bio-Rad, USA).

### Zeta Potential Analysis

The electrokinetic potential of exosomal suspension was measured to check for particle stability in suspension. Samples were diluted in 1:3000 ratio and were loaded into the disposable capillary cells, Malvern Panalytical (DTS1070) zeta potential analyser. The magnitudes were read in triplicates and the average spectra were considered for comparative purpose.

### Raman Spectroscopy (Labram HR Evolution - Horiba)

Labram HR Evolution – Horiba Micro Raman Spectroscopy (Kyoto, Japan) was used for the spectral recording of exosomes for label-free molecular characterization using a 532nm laser at 25% laser intensity. The molecular fingerprint range and high-frequency range of Raman shift used for the study were 500 cm^−1^ to 1800 cm^−1^ and 2600 cm^−1^ to 3200 cm^−1^. The sample was made to airdrop on a glass slide and air-dried overnight with a desiccant in a closed chamber. Brain sphingomyelin (SM) and Cholesterol (CHL) purchased from Avanti Polar Lipids, (Alabaster, AL, USA) were used to acquire reference spectra. The acquisition times used for recording spectra for control and the samples were 10 and 15 respectively.

### Fourier – transform infrared spectroscopy

ATR-FTIR IR Tracer 100-Shimadzu (Kyoto, Japan) was used for the spectral recording of exosome samples. The samples were dried to powder form for label-free molecular characterization. The ATR-FTIR spectral recordings were considered from the wavelength range of 500 cm^−1^ to 1800 cm^−1^ and 2600 cm^−1^ to 4000 cm^−1^, the molecular fingerprint range and high-frequency range. Samples were experimented along with the reference samples of brain sphingomyelin (SM) lipid and Cholesterol (CHL) purchased from Avanti Polar Lipids, (Alabaster, AL, USA) each with acquisition parameters of 50 scans and 4 cm^−1^ resolution. Normalization and multipoint baseline correction of the spectra were done and Happ - Genzel function and Savitzky-Golay algorithm were used for apodization and smoothing of the spectra at the beginning and end of the time-based sampling using the LabSolutions IR.

### Data analysis

Multipoint baseline correction, normalization, and smoothing of the Raman and ATR-FTIR spectra were done in Lab Spec 6 and LabSolutions IR respectively with the given parameters. After the data acquisition, the digital numeric values having wavelength and intensity values on X axis and Y axis respectively were plotted simultaneously for all the samples along with the reference spectra. For prediction and determination of the biological components from the spectra based on their molecular vibrations, curve fitting and deconvolution were performed in OriginLab Pro 2020 ver. 9.4.9. Chi^2^ and R^2^ followed by regression analysis of the curves were optimized by performing 400 iterations of non – linear regression analysis performed using the Levenberg Marquardt algorithm with Lorentz function to provide best fit curves and assessment of ease of determination of molecular components.

Protein to lipid ratios for the three techniques were determined from the integrated area of regions of interest (ROI) corresponding to protein (1400 – 1800 cm^−1^)and lipids (2700 – 3040 cm^−1^) in RAMAN and ATR-FTIR spectra. The area under curves were calculated by the integral function calculating mathematical area with limits fit to interpolate the region of curves having baseline mode Y = 0 following which the ratios or relative intensities were determined by the division of these integrated areas of proteins and lipids of individual spectral recordings (supplementary table S3).

Statistical analyses were done for understanding the variation in spectral components of the Raman and ATR-FTIR spectra obtained for the exosomes isolated from two cell lines by three different isolation techniques. The tests were done in Origin Lab Pro 2020 ver. 9.4.9. Levene’s test was performed to check for the equality of variances in the spectral data. Following this One-way ANOVA and Tukey’s test were performed to check for the statistical evidence of significantly different population means and to find the variable responsible for significantly different means in population with the actual power of 0.05 significance level.

Multivariate analyses in the form of Principal component analysis (PCA) of the normalized spectra were performed to reduce the dimensions or the variability in the data and determine their principal components (PCs) or major trends that vary the most from the mixed population. Grouped PCA on principal component scores was performed to establish discrimination in the variation of data and verification of the difference in their mean, irrespective of data being within-group variance.

## 3. Results and Discussions

### Quantification of the exosome yield and exosome Proteome

The Exosome samples isolated from COLO 205 and MCF 7 were resuspended in 50 μl of NFW (for imaging purpose) and 1 X PBS and stored in −20°C freezer for further protein analysis. For quantification, the samples were thawed on ice and were used for protein quantification. Exosome quantification and exosomal total protein quantification (figure 3.1 and 3.2)

**Figure 3.1.**
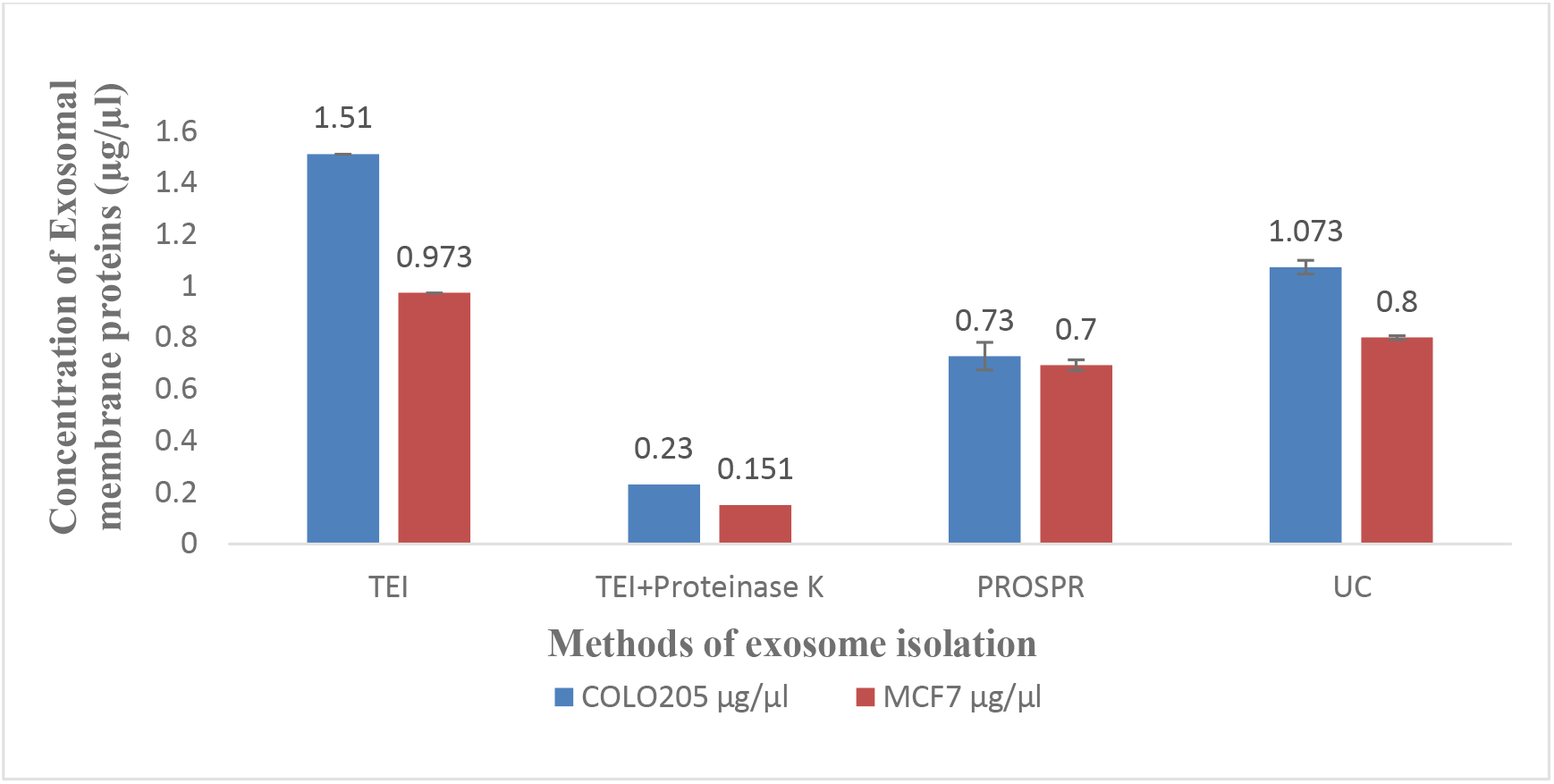
Bar graph representing the concentration of exosome yield per 10^6^ cells obtained from COLO 205 and MCF-7 exosomes isolated by the three techniques. Blue and orange bars represent COLO 205 exosomes and MCF-7 exosomes concentration respectively.

**Figure 3.2.**
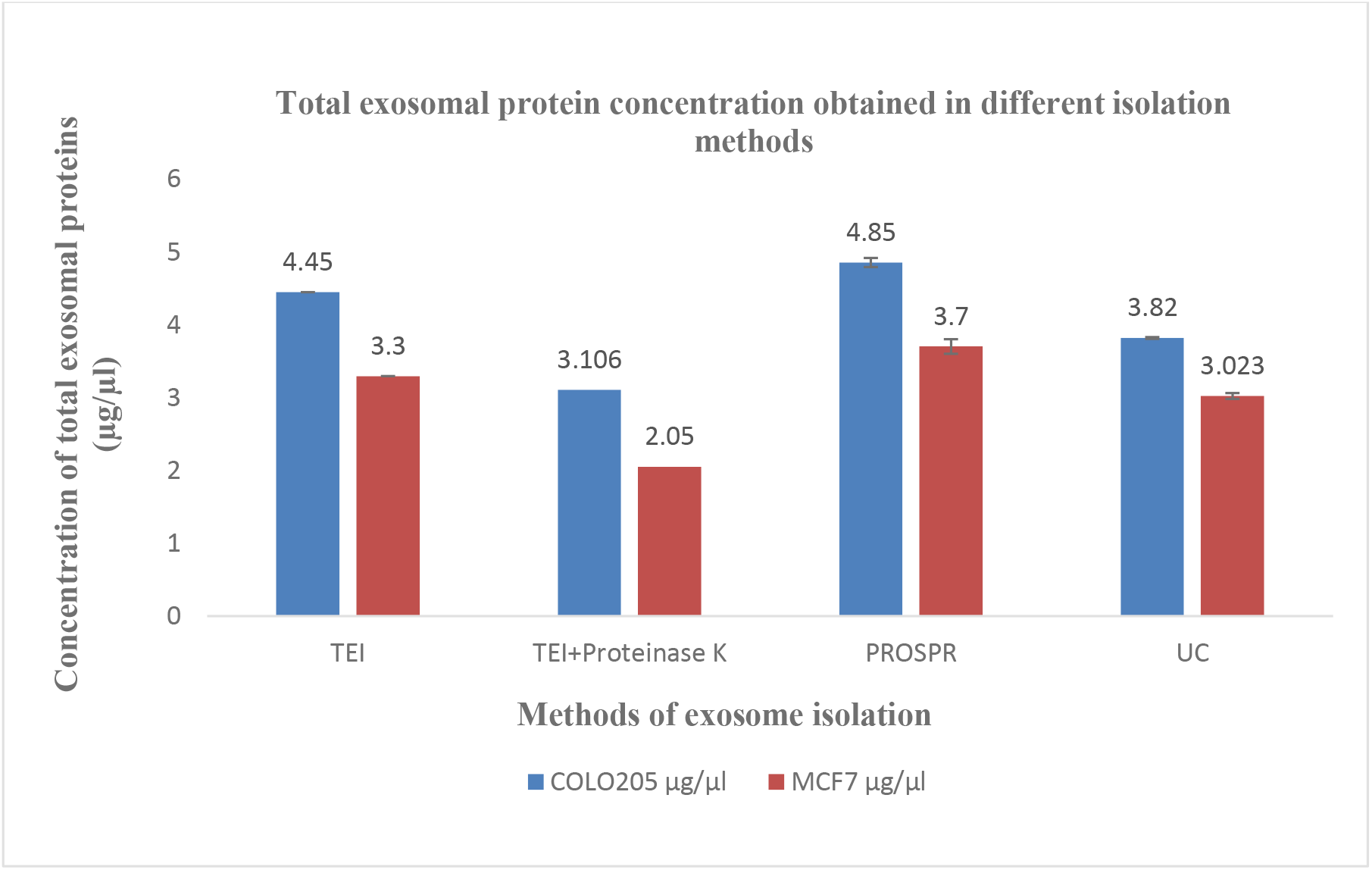
Bar graph representing the concentration of exosomal protein concentration per 10^6^ cells obtained from COLO 205 and MCF-7 exosomes isolated by the three techniques. Blue and orange bars represent COLO 205 exosomes and MCF-7 exosome concentration respectively.

Quantification of exosomes using Bradford’s assay reading the absorbance at 595nm is an indirect quantification approach. The protocol uses Bradford’s reagent that binds to the proteins, and indirectly indicate the number of exosomes by quantifying the surface proteins and receptors^43^.

Yield from PROSPR method was less in terms of exosome quantification but high in total protein quantification when compared with TEI and UC. TEI on the other hand yielded more exosomes and higher protein content compared to UC (figure 3.2). The proteinase K treated samples showed drastic reduction in the exosome count as the proteinase K is probably degrading the surface proteins present in the exosome membrane and hence can be useful for analyzing the exosome cargo but not for enrichment of purity. In case of exosome’s hydrodynamic diameter, the treatments like trypsin or proteinase K digestion have also been shown to wear off the electric dipole layer adhering to the surface proteins on membrane due to their digestion^45^. This will affect the morphology of exosomes isolated.

### Characterization of Exosomes

#### High-Resolution Transmission Electron Microscopy

The HRTEM images of exosomes confirmed their typical morphology with the lipid bilayer membrane and were further used for image analysis to compare the morphological attributes of exosomes and their particle size distribution. Our simple procedure of sample preparation resulted in high quality images comparable to a negative staining protocol. The exosomes isolated from COLO 205 and MCF-7 conditioned media showed distinct morphologies for each case when isolated from two different sources with three different techniques. The exosomes isolated from the two cell lines by the three different techniques are presented in figure 3.3.

**Figure 3.3.**
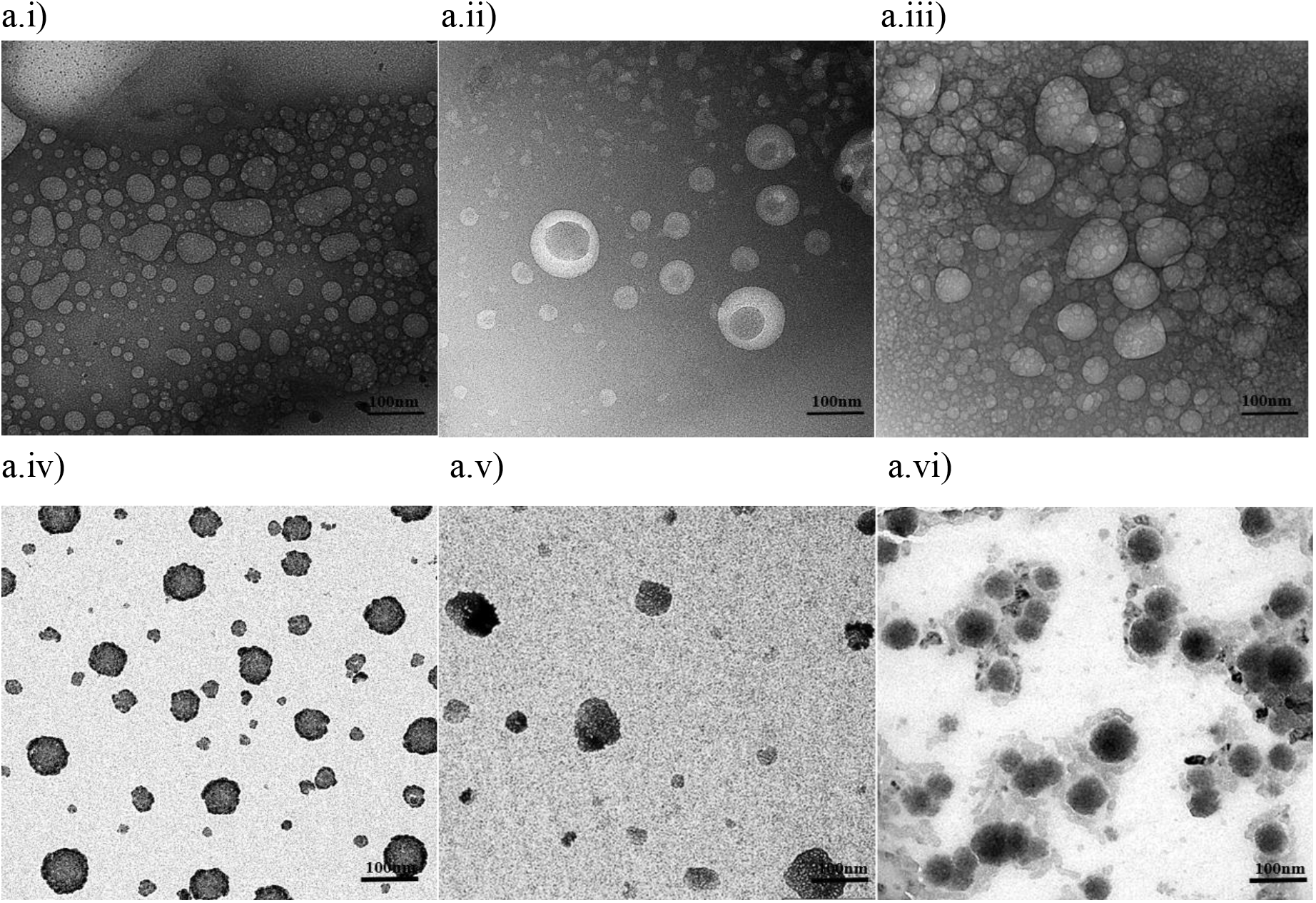
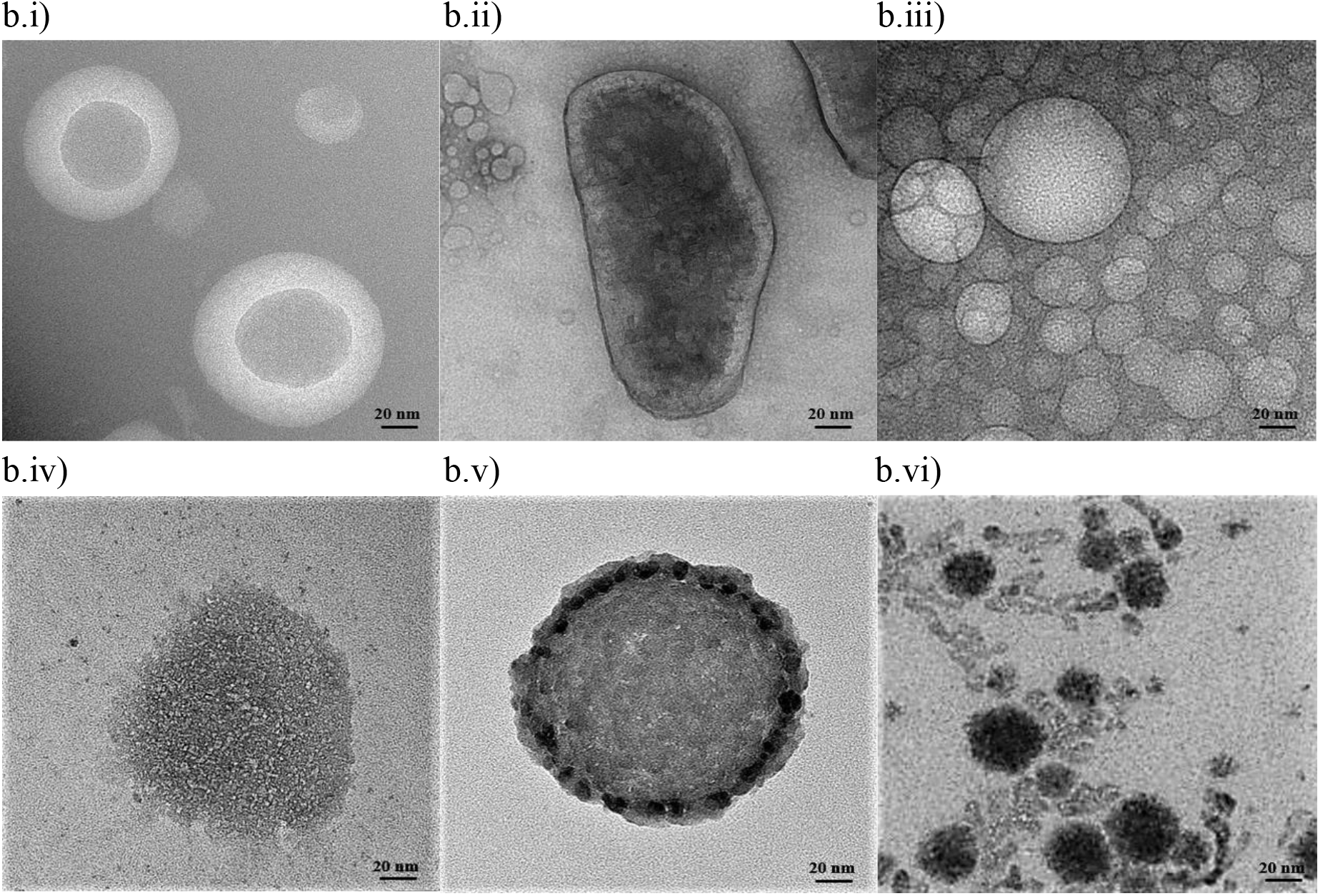
High-resolution transmission electron microscopy of exosomes isolated from COLO 205 and MCF-7 media. I) COLO 205 exosomes isolated by TEI, ii) COLO 205 exosomes isolated by PROSPR, iii) COLO 205 exosomes isolated by UC, iv) MCF-7 exosomes isolated by TEI, v) MCF-7 exosomes isolated by PROSPR and vi) MCF-7 exosomes isolated by UC respectively. The images presented have a resolution of 3.3 a) 100nm and 4b)20 nm.

### Image Analysis of the HRTEM images obtained for Exosomes

#### Interactive surface 3D plots for Morphology Study

The optimised parameters in the interactive environment of analysis manager of FIJI ImageJ were used to plot high resolution morphology surface plots, thermal surface plots and distribution surface plots (figure 3.4). The images and analysis of the morphology interpret differential morphology of exosomes when isolated via the three different techniques and two different cell sources.

**Figure 3.4.**
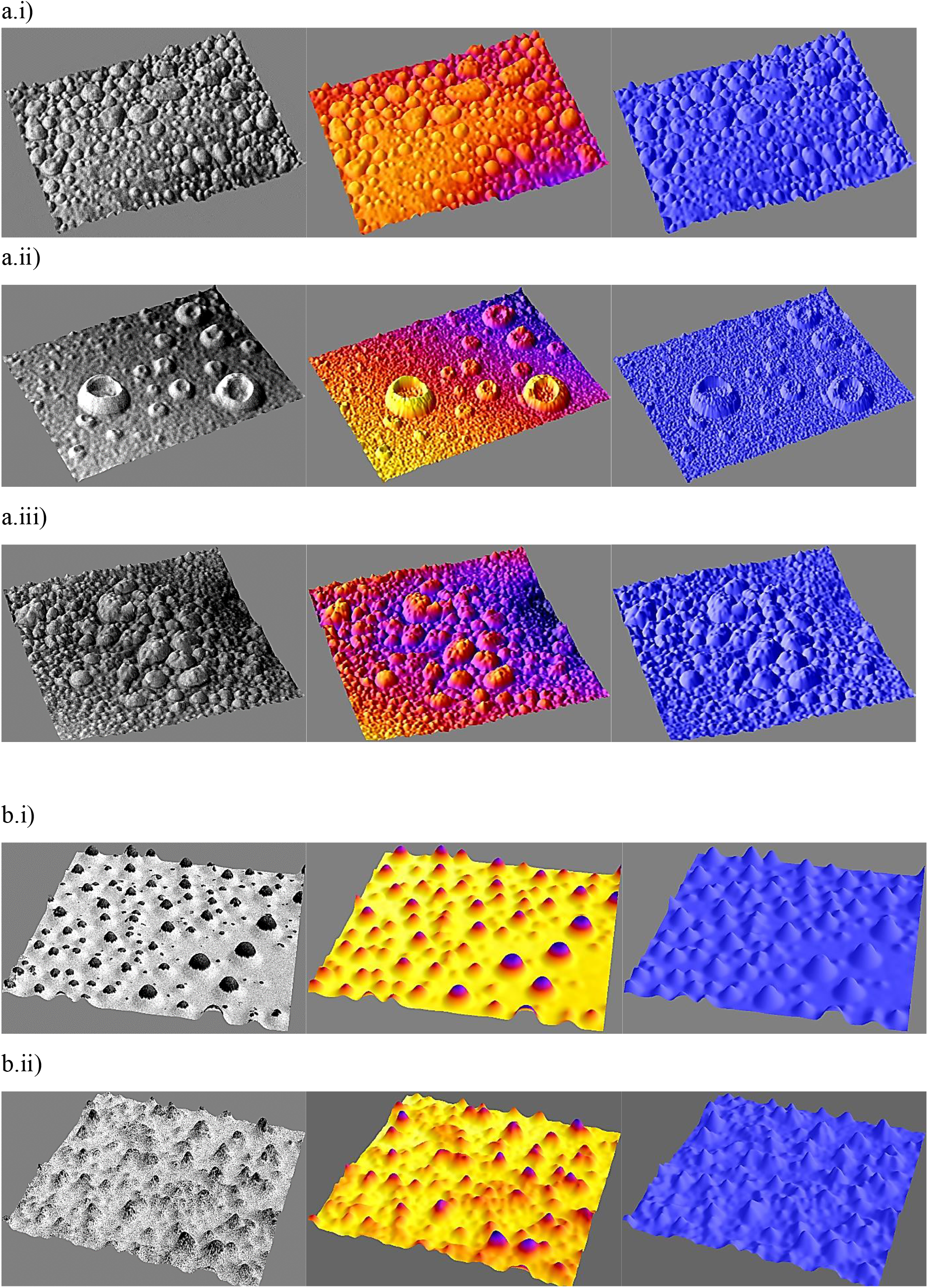
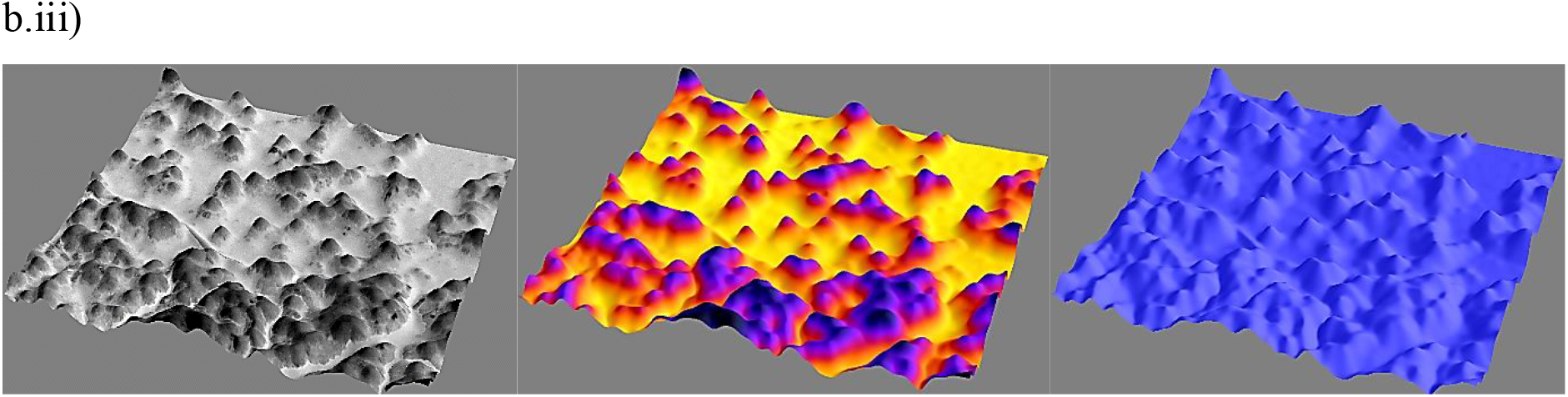
3D high resolution interactive surface plots plotted for morphology study of 9a) COLO205 and 9b) MCF-7 exosomes. i) Topology map plotted for the study of exosome morphology. ii) Thermal map plotted for distinguishing between exosomes and noise. iii) Distribution map plotted for studying the dispersion and distribution pattern of exosomes.

Morphology surface plots or topology plots presented the 3D morphology of exosomes and their physical attributes. Thermal plots depicted the difference between true exosome vesicles and background noise for size distribution analysis. Thermographs with thermal signature of exosomes rising to peaks were considered true exosomes and were used for the calculation and statistics. Distribution plots depicted the pattern of dispersion and distribution of exosomes in suspension also gave insight into the pattern of distribution of small and big exosomes.

The exosomes isolated by TEI and UC were of oval shaped as reported in literature^46–48^. But in the case of PROSPR change of morphology was observed. The exosomes isolated from TEI showed better recovery of morphology and well dispersed reflecting towards better stability of particles. The PROSPR technique yielded bulged and cup-shaped, not much globular exosomes and other bigger microvesicles. from both COLO 205 and MCF-7. In the case of exosomes isolated by UC, the COLO 205 and MCF-7 exosomes both showed particle clustering and agglomeration in various regions along with background noise. This noise was supposedly of aggregated proteins and other small artifacts around these exosomes that might be responsible for the particle aggregation^49^. It was already reported that in urinary exosomes, proteins called Tamm-Horsfall induced exosome aggregation when isolated by ultracentrifugation. These proteins comes down along with exosomes down due to high centrifugal force^50,51^.

### Plotting 2D contour map for Particle Size Distribution Analysis

HRTEM images provided accurate information about the size and morphology of the particles. Calculating the Sauter mean diameter or the surface - volume mean diameter and Feret’s diameter or caliper diameter of exosomes based on their absolute morphology provided the accurate information of their sizes in sample holding an advantage over Dynamic Light Scattering^22,48,52–55^. 2D contour maps were plotted in Origin Lab and the parameters were recorded. (Supplementary figure S1). The data was obtained from uniformly scaled and calibrated images.

### Normal Distribution curves, Particle size frequency distribution curves and comparative box plot for particle size distribution

Normal distribution curves for particle area, volume and size were plotted for both COLO 205 and MCF-7 exosomes based on these data (Supplementary figure S2, S3 & S4). The parameters were analysed based on the standard deviation of area under the curve with a set threshold. Curves for exosomes isolated by TEI had even distribution with a broader size range compared to UC which had steepest curves among the three with narrow size range. PROSPR surpassed the threshold in all the distribution curves in both cases of COLO205 and MCF-7 exosomes. Based on this data, particle size frequency curves (Supplementary figure S5) and frequency distribution curves (Supplementary figure S6) were plotted. Higher percentage of exosomes occurred between 30-50 nm in all three techniques. Comparative box (figure 3.5) plots were plotted based on the quartile deviations in the data (Supplementary Tables S1 & S2) containing particle count and size which validated the above observation giving clear information about mean size, range and outliers. In case of TEI the exosomes had the mean size of 40-50nm with max ranging to 190 nm, in UC mean size was 30 nm and max was 170nm. The PROSPPR samples had the average mean size of 40-60nm with outliers reaching up to 600nm and above that indicated the presence of bigger microvesicles in both COLO205 and MCF-7 exosomes.

**Figure 3.5.**
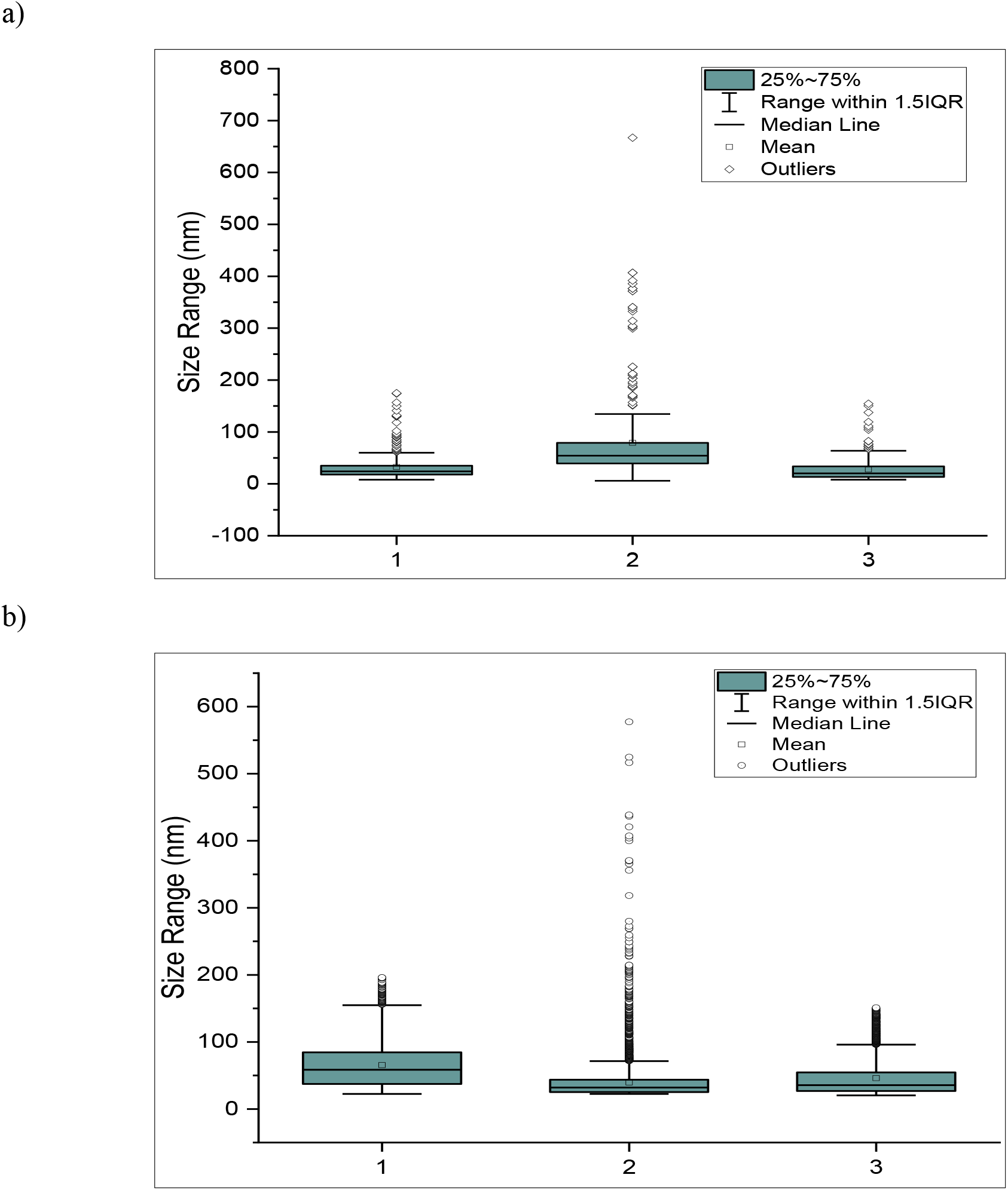
Comparative box plots of particle size distribution for exosomes isolated from COLO 205 and MCF-7 by TEI, PROSPR and UC techniques. The box plots were plotted based on the quartile deviation in data. The box plots comprised of the mean size and 25-75% data within 1.5 interquartile range. Outliers in the data lied outside the 1.5 IQR that helped determine the size of exosomes frequently occurring and the ones significantly differing from other data points. a) Comparative box plots of COLO 205 exosomes and b) comparative box plots of MCF-7 exosomes. The numbers 1,2 and 3 labelled on x – axis in figure 3.5. a) and b) represent TEI, PROSPR and UC respectively.

### Zeta Potential analysis

The electrokinetic potential test helps assess the stability of exosomes in terms of mV ranging from incipient stability to good stability in suspension. Exosomes isolated by TEI had zeta potential magnitude above −50 mV resulting in good stability of particles with very less agglomeration. The PROSPR and UC processed samples had zeta potentials in the range −10 to −40 mV resulting in moderate and incipient stability of particles. The UC samples had the lowest zeta potential values than others that suggested exosomes in suspension might be facing aggregation or agglomeration (figure 3.6). The individual zeta potential graphs can be found in supplementary data (Supplementary figure S7).

**Figure 3.6.**
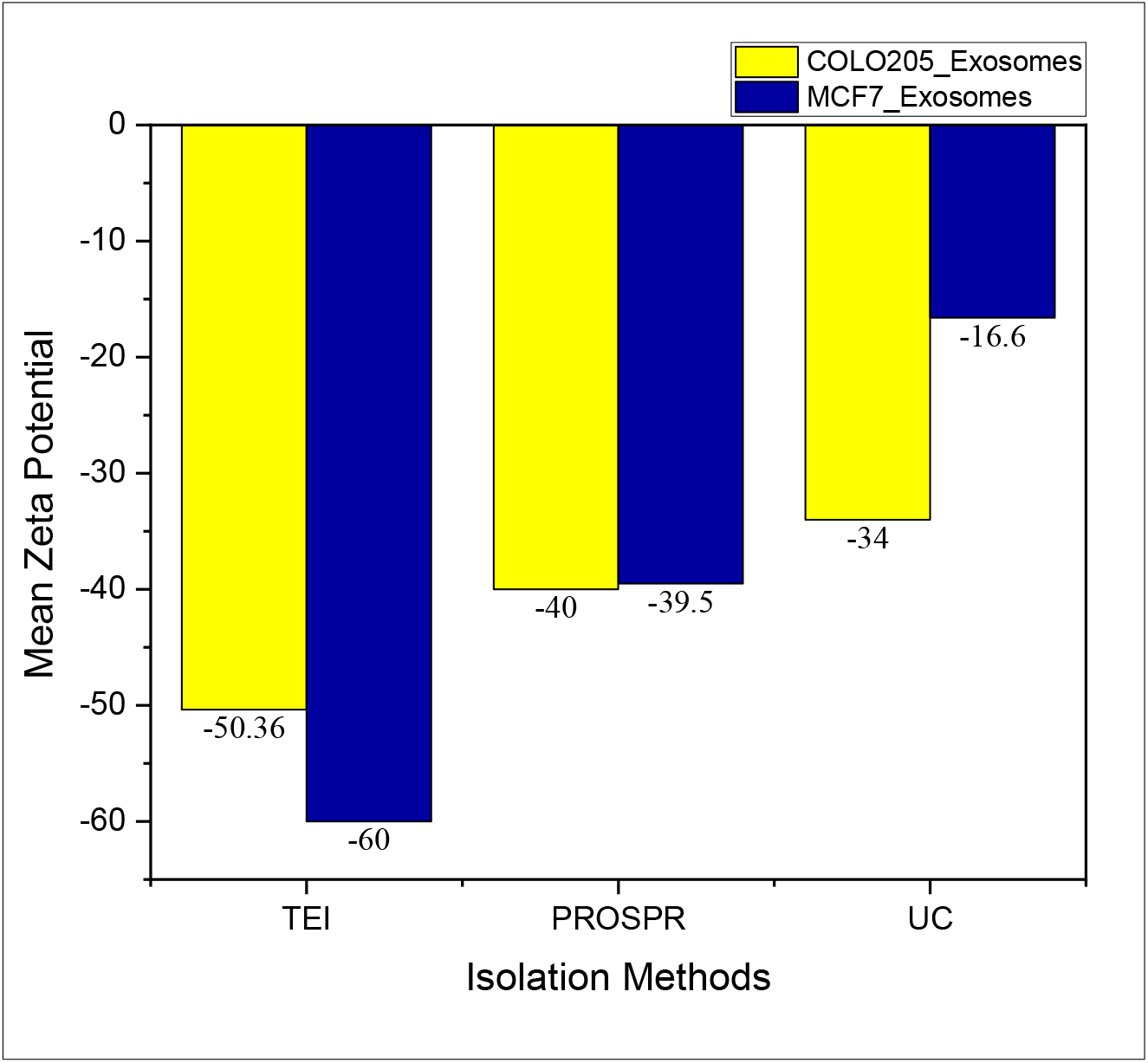
Comparative zeta potential analysis: The stability of exosomes in suspension was analysed by their electrokinetic potential. The yellow bars represent COLO 205 exosomes and blue bars represent MCF-7 exosomes.

### SDS PAGE of total protein in exosomes and western blotting for immuno-characterization

Exosome samples having protein concentration of ~ 150 μg to 200 μg in 100 μl of suspension were loaded into PAGE (figure 3.7 & Supplementary figure S8). Many peptides that were visible in TEI and UC based isolation were absent in PROSPR based isolation method. TEI and UC based isolation methods yielded peptides in the range of 10 kDa to 250 kDa where TEI method had better band intensities compared to others. In case of PROSPR, only 55kDa to 70kDa peptides were visible making it the method with least protein recovery. The results stay the same for both the cell line sources.

**Figure 3.7.**
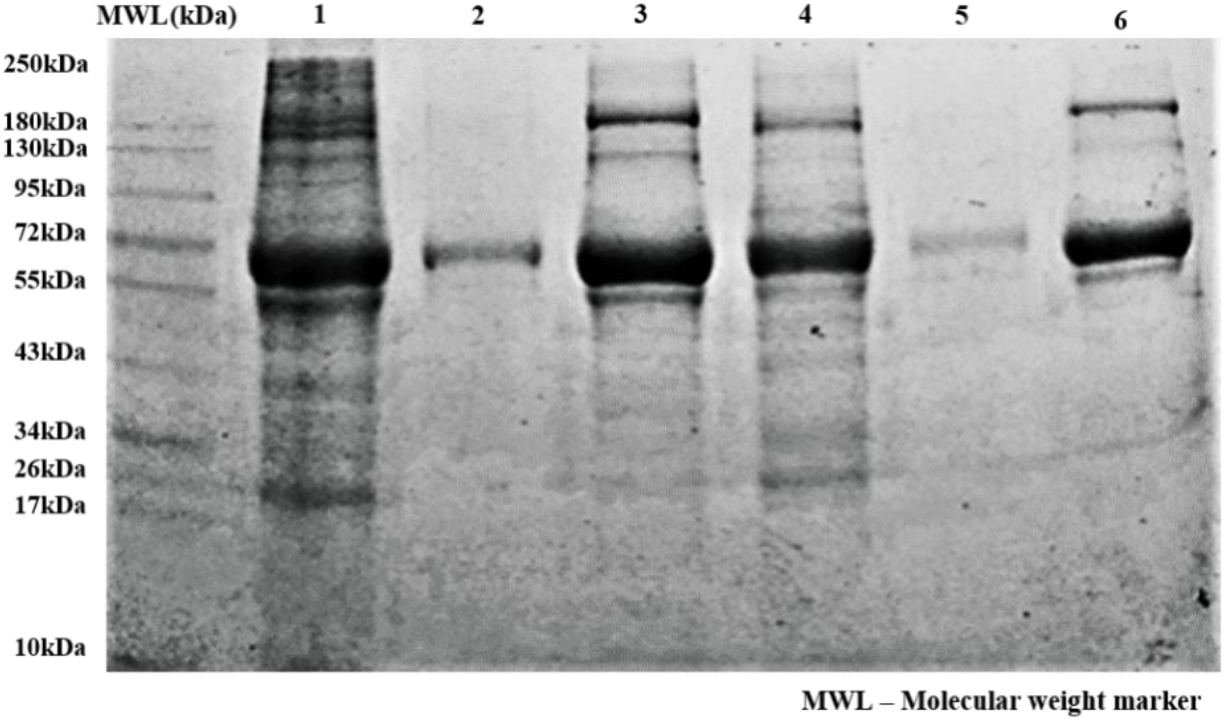
SDS PAGE for the exosome proteome profile. The 10% separating gel was used to run the sample in gel based on their charge and mass. 1^st^ lane was the molecular marker with the range of 10-250 kDa, 2^nd^ lane onwards samples loaded were COLO 205 TEI, PROSPR, UC and MCF-7 TEI, PROSPR, UC samples consecutively.

Western blotting by targeting CD63 differentiated the techniques based on protein recovery. The order of intensities was TEI > UC > PROSPR showing highest recovery of CD63 in TEI versus UC and least in PROSPR (figure 3.8).

**Figure 3.8.**
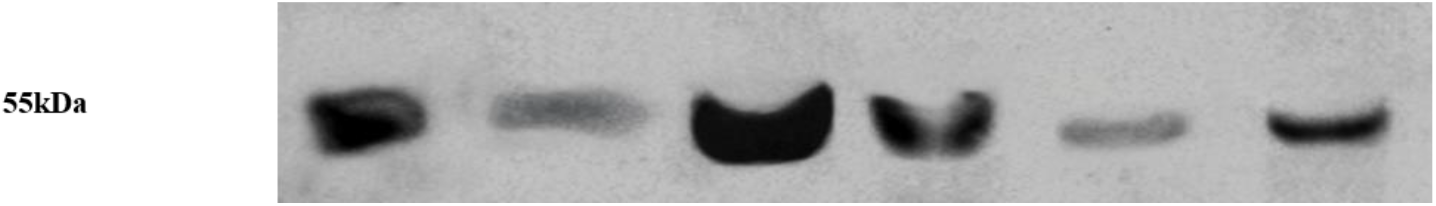
Western Blotting of exosomes for molecular characterization with CD63 marker. The samples blotted were COLO 205 UC, COLO 205 PROSPR, COLO 205 TEI and MCF-7 UC, MCF-7 PROSPR, MCF-7 TEI consecutively from the first lane.

### Molecular and analytical characterization of exosomes by label free fingerprinting with micro Raman Spectroscopy and ATR-FTIR

RAMAN and infrared spectroscopic approaches provide the advantage of studying various molecular constituents of exosomes through the vibrational conformation of their atoms and molecules present in structural backbone or functional groups^56,57^. These approaches were used to study the molecular dynamic characteristics of exosomes, that required very less sample (<1mM) for the analysis. ATR-FTIR spectroscopy provided much accurate spectra but couldn’t provide information on macromolecular components that RAMAN spectroscopy provided. ATR-FTIR provided spectra without the problem of light scattering and background noise or fluorescence. In case of ATR-FTIR, the absorption of water molecules was subtracted from the background^58,59^. The techniques have been employed in recent studies to identify the molecular components of exosomes in a label free manner, differentiating them in disease versus non diseased state and discriminating the exosomes from other vesicles. We thought to employ the same principle of label free characterization of exosomes to differentiate the techniques and categorize them based on ease of performing the experiment and determining the molecular components of exosomes, quality of spectra obtained suggesting a technique with least deviations. Refer the supplementary material for the schematics of regression models, Chi sq and R^2^ values and the exosomal molecular components determined from the spectra of both techniques (Supplementary figure S9, supplementary table S3 & S4). The RAMAN and FTIR spectra for exosomes isolated from COLO 205 and MCF-7 by TEI, PROSPR, and UC are presented in figure 3.9.

**Figure 3.9.**
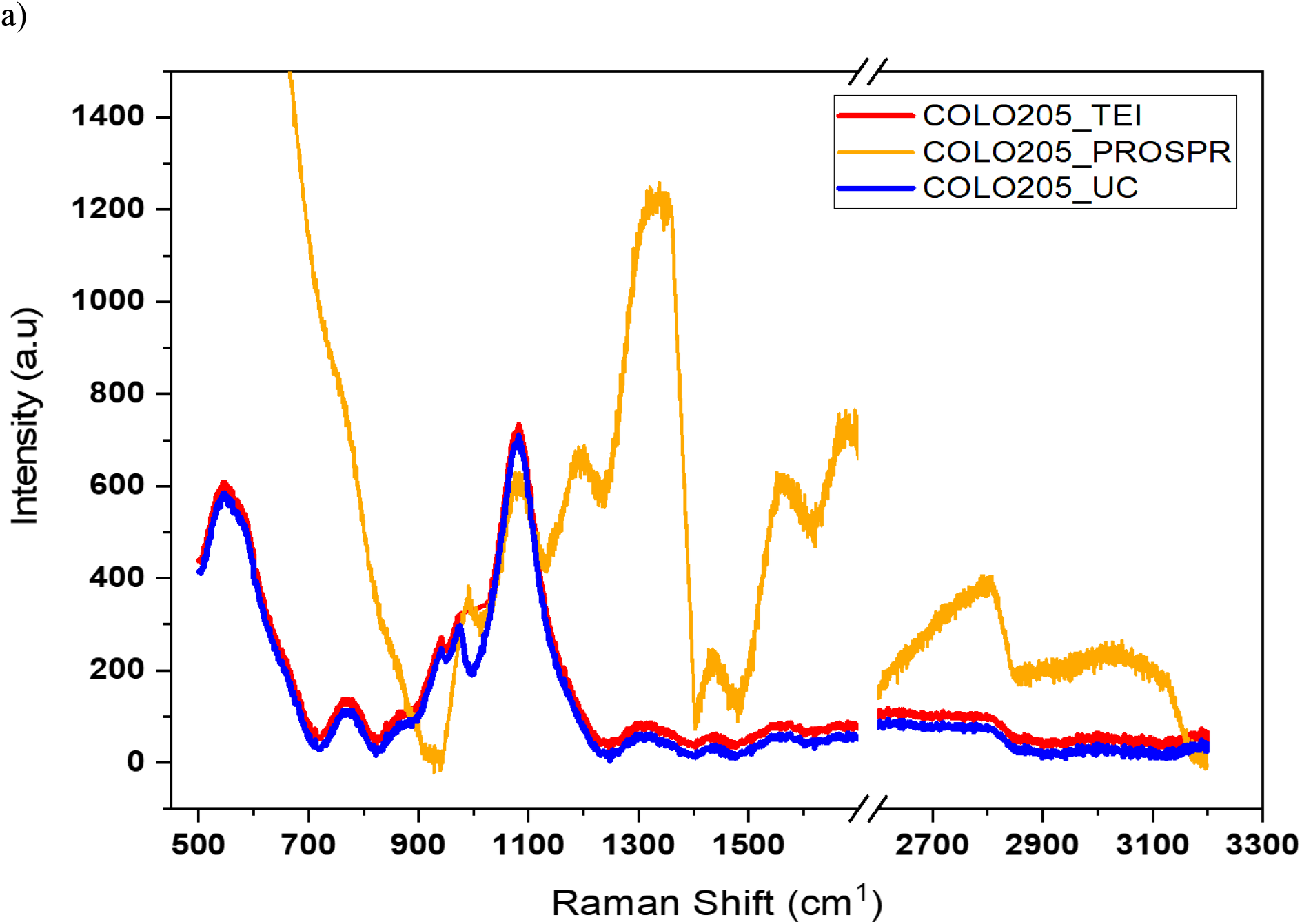
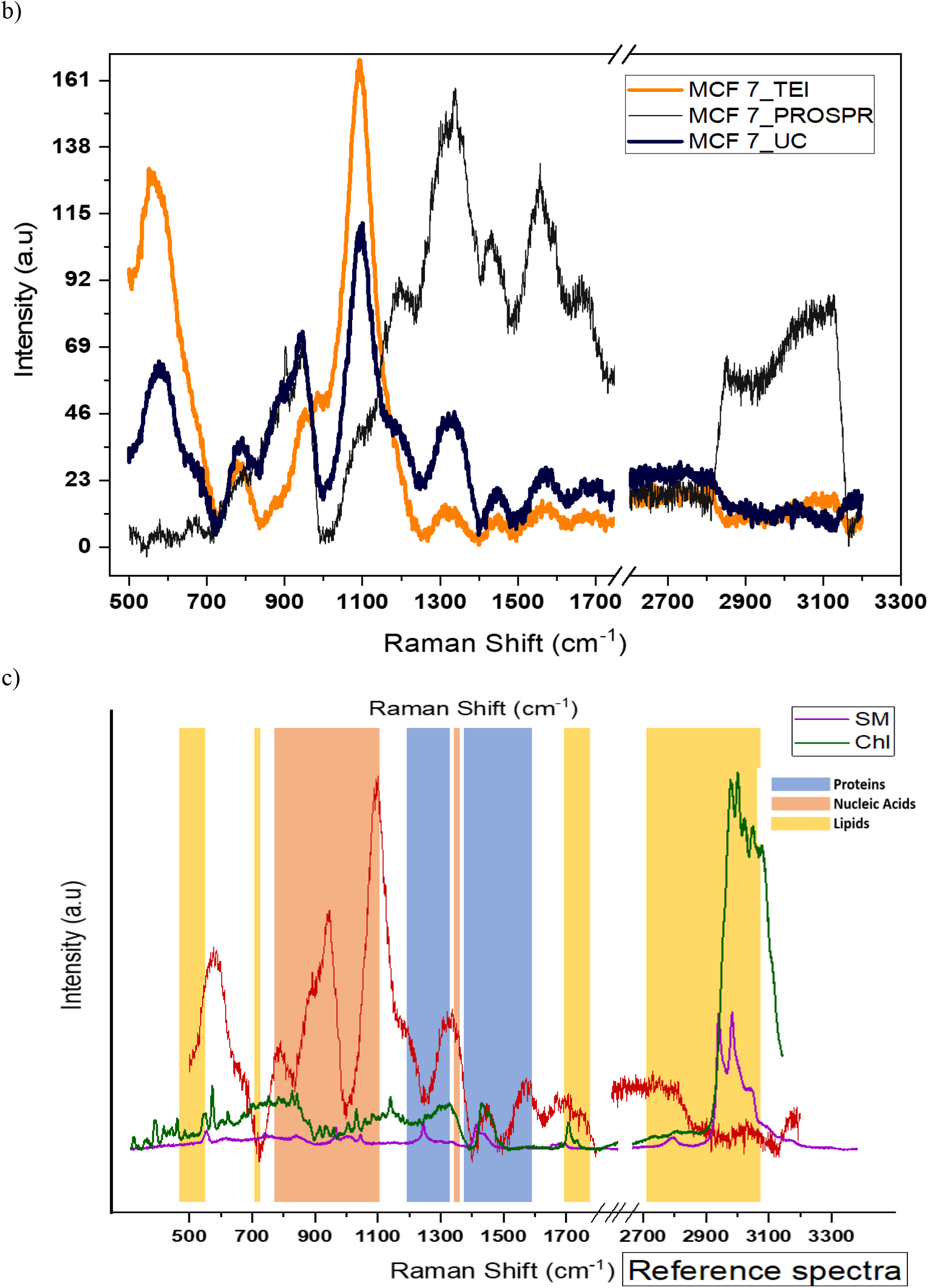
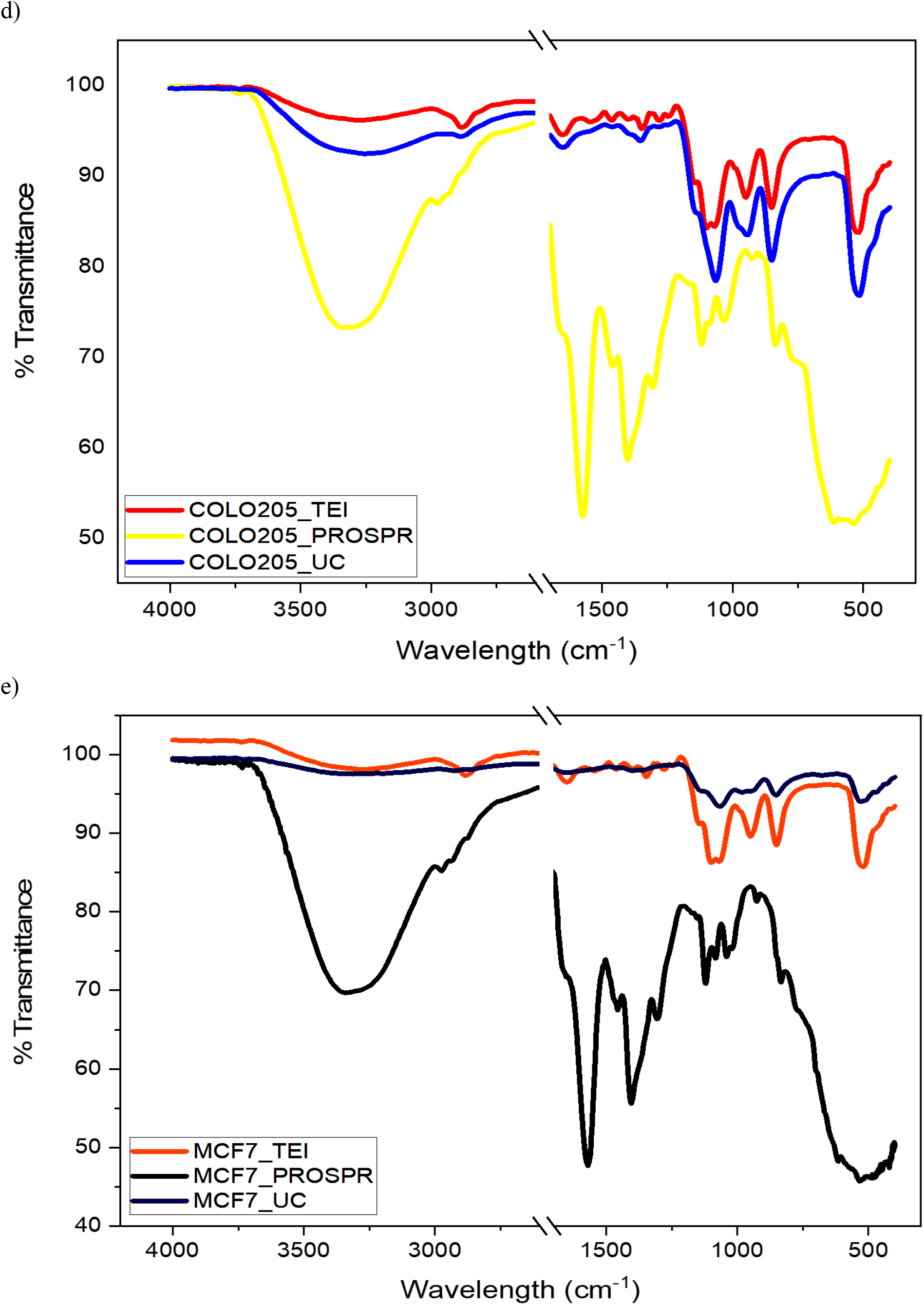
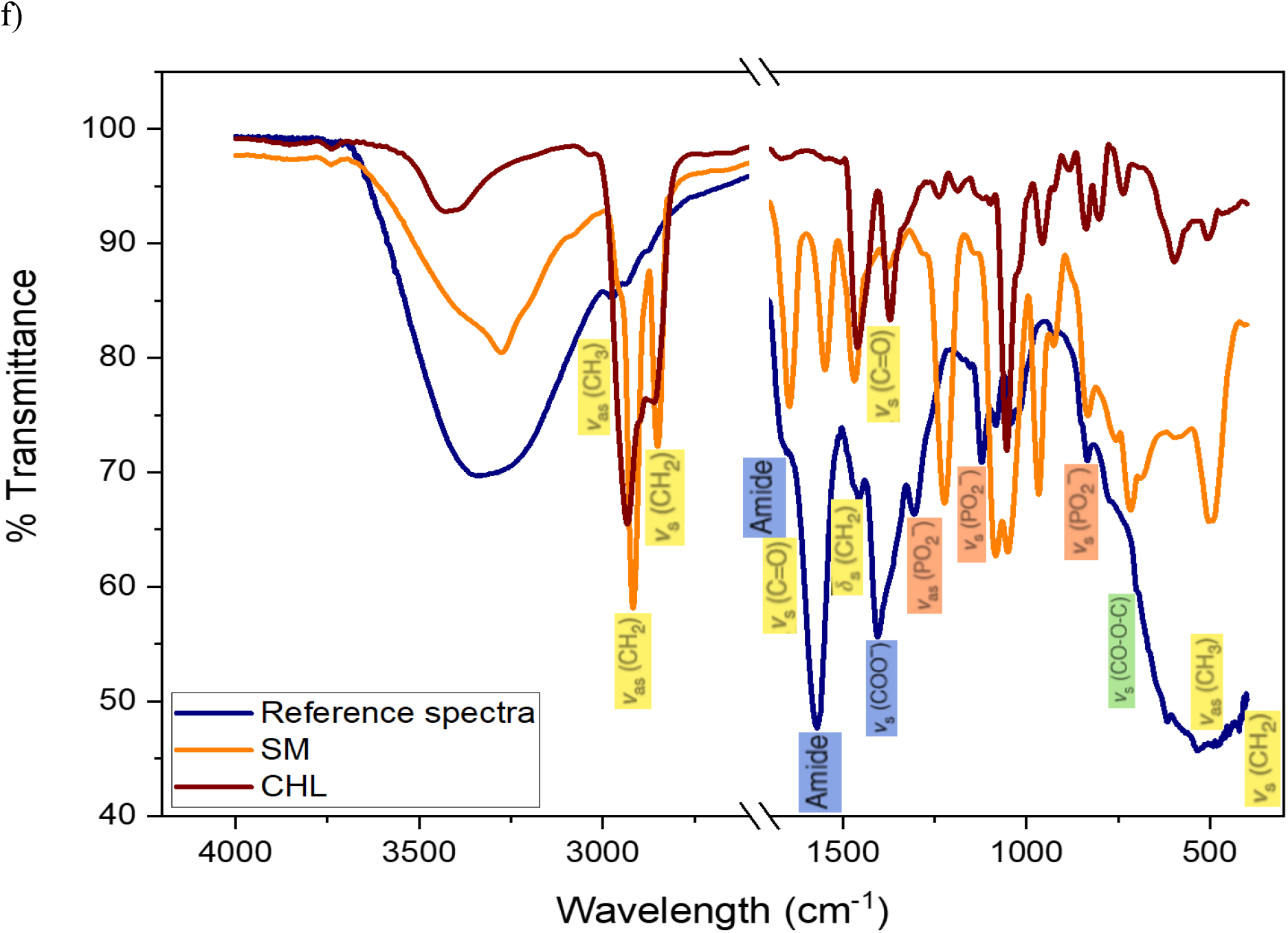
a) RAMAN spectral presentation of COLO 205 exosomes isolated by the three techniques, TEI (green), PROSPR (violet), and UC (orange). b) RAMAN spectral presentation of MCF-7 exosomes isolated by the three techniques, TEI (black), PROSPR (red) and UC (blue). c) Reference spectra for RAMAN spectroscopy depicting the potential regions of biomolecules representing the cargo and structural compositions of exosomes. labelling of the regions with blue represent protein species, orange regions represent the nucleic acids species and yellow regions represent the lipids species in exosomes. d) ATR-FTIR spectral presentation of COLO 205 exosomes isolated by the three techniques, TEI (green), PROSPR (violet), and UC (orange). e) ATR-FTIR spectral presentation of MCF-7 exosomes isolated by the three techniques, TEI (black), PROSPR (red) and UC (blue). f) Reference spectra for ATR-FTIR spectroscopy depicting the potential regions of biomolecules representing the cargo and structural compositions of exosomes. Molecular representation in yellow labels represent species of lipids with symmetric and asymmetric vibrations of CH_2_ and CH_3_, green labels represent species of carbohydrates with vibrations of carboxyl group, orange labels represent species of nucleic acids with vibrations of phosphate group and blue labels represent species of proteins with vibration of amides in exosomes.

During the analysis some interesting observations were definitely noted while deconvolution of peaks in the 1400 – 1800 cm^−1^ region. One main observation was the presence of spectral intensities at 1622 cm-1 corresponding for aggregated proteins or apolipoproteins in the UC and PROSPR exosomal spectra but the same was a flattened curve in the case of TEI exosomes of both the cell lines. This supported the exosome distribution and zeta potential analysis.

#### Spectroscopic protein to lipid ratio

The relative intensities were plotted on to bar plots for both COLO 205 and MCF-7 exosomes isolated by TEI, PROSPR and UC (figure 3.10) where the order of the ratios was PROSPR > TEI > UC. The relative intensity of TEI was slightly higher than UC which correlated with the total protein estimation and exosomal yield but the greater ratio in PROSPR exosomes directly correlated to the total protein estimation. The observation was validated with one-way ANOVA and Tukey’s test which had non-significant mean difference in TEI-UC combination but significant mean difference in TEI-PROSPR and UC-PROSPR combinations at 0.05 level (Supplementary figure S10 for Mean comparison plot and Supplementary table S5 for spectroscopic protein to lipid ratios).

**Figure 3.10.**
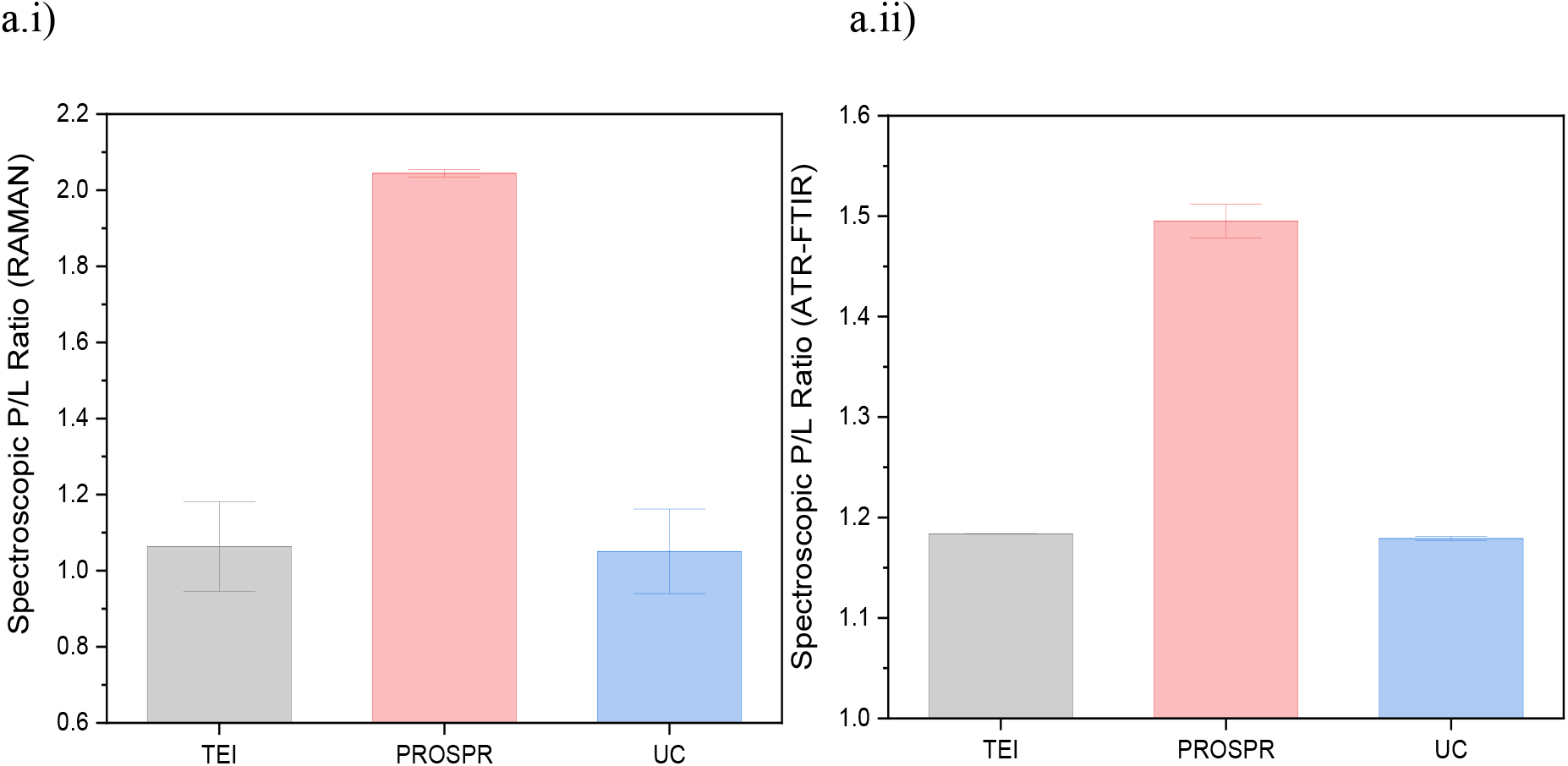
Representation of relative intensities for spectroscopic protein to lipid ratio determined from a.i) RAMAN spectra and b.i) ATR-FTIR spectra by division of integral areas (IA) calculated for proteins and lipids (IA (1400 – 1800 cm^−1^) / IA (2700 – 3040 cm^−1^)) of COLO 205 and MCF-7 exosomes isolated by TEI, PROSPR, and UC. The data was statistically analysed and validated with one-way ANOVA and by comparison of means of the relative intensities of a.ii) spectroscopic P/L ratio of RAMAN spectra and b.ii) spectroscopic P/L ratio of ATR-FTIR spectra

#### Validation of data with statistical analysis

Statistical analysis for spectral data obtained from both the analytical techniques showed similar trend in results for COLO 205 and MCF-7 exosomes isolated by three different isolation techniques. The results of descriptive statistics reflected towards the presence of large amount of variation in the spectral data. Samples isolated from TEI and UC had similar but smaller standard deviations from mean as compared to the PROSPR samples. Levene’s test proved that the population variances were significantly different at 0.05 significance level having the power as 1 rejecting the null hypothesis. The one-way ANOVA performed showed population means to be significantly different which validated the acceptance for alternate hypothesis at 0.05 levels. Following one-way ANOVA, Tukey’s test resulted in TEI and UC methods in combination to have non-significant mean difference at 0.05 levels but the other combinations of TEI-PROSPR and UC-PROSPR had significant mean differences at 0.05 levels (Refer to Supplementary tables S6–S9 for statistics data and supplementary figure S11 for the Mean comparison plots).

Grouped PCAs were plotted (figure 3.11) with two principal components PC1 and PC2 having 57.5% and 42.4% variance in RAMAN spectra and 99.84% and 0.15% variance in ATR-FTIR spectra (Scree plots in Supplementary figure S12 & S15). The plots discriminated the TEI, UC, and PROSPR samples based on component scores and COLO 205 and MCF-7 exosomes based on the vectors of loadings. The vectors were not completely out of phase by 90° but were separated with a greater phase angle that differentiated exosomes based on their sources. Component scores of TEI and UC data showed tight clustering due to strong correlation but PROSPR that had correlation with TEI and UC also had more variables distributed with its confidence ellipse interpreting larger group differences in the canonical space.

**Figure 3.11.**
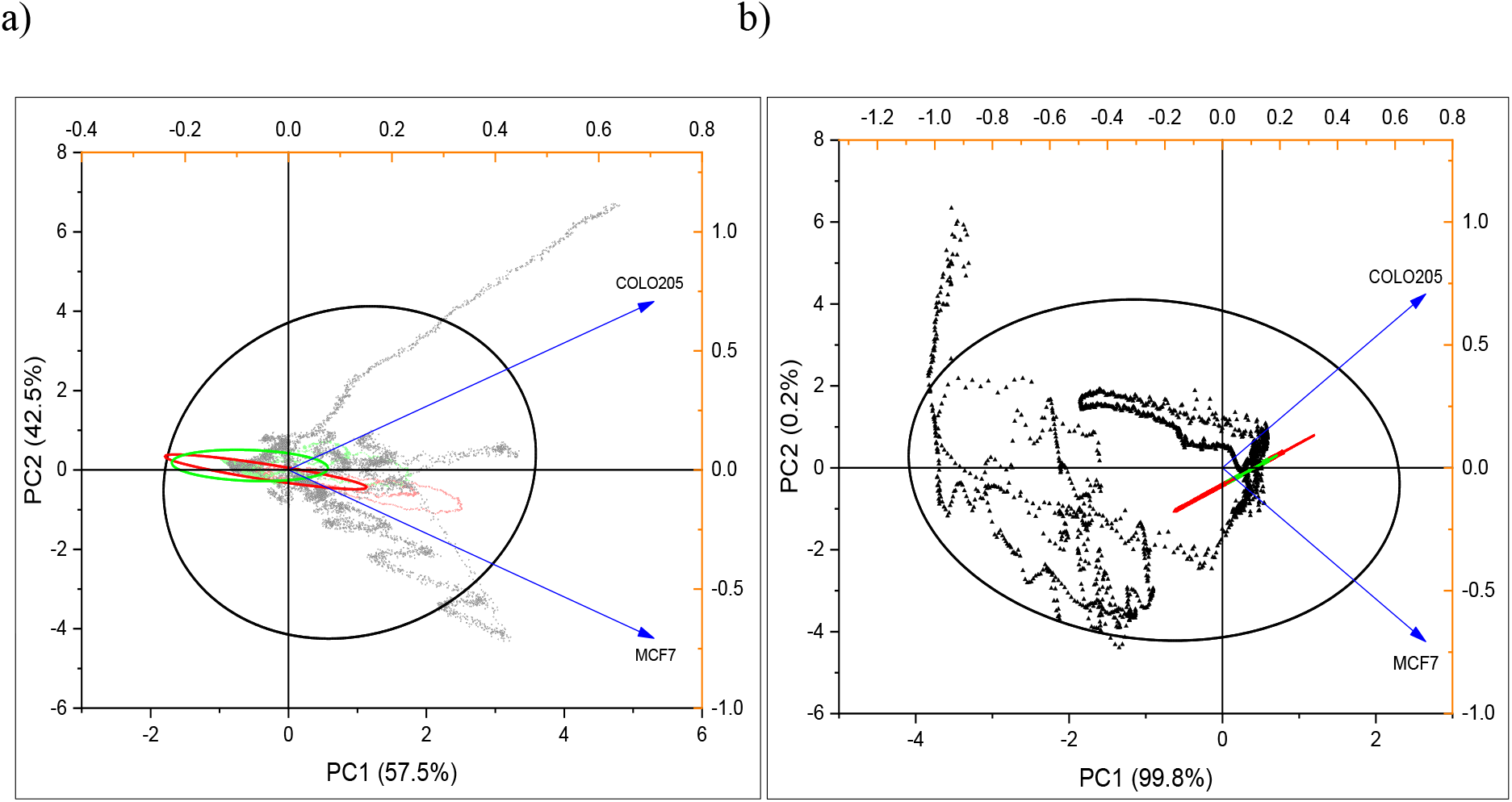
Grouped PCA plot for a) RAMAN spectra and b) ATR-FTIR spectra recorded for COLO 205 and MCF-7 exosomes isolated by TEI, PROSPR, and UC. PCA loadings in blue represent the exosome types and scores represent the grouped spectral data for TEI (red), PROSPR (black) and UC (green). Confidence ellipses in red represent TEI, in black represent PROSPR and in green represent UC.

In case of COLO 205 and MCF-7 exosome types individually, PCA was performed with two principal components for both spectroscopic techniques with loading vectors designated as isolation techniques to identify the phase difference between each of them. This was presented in biplots (Supplementary figures for biplots S18 and scree plots S13, S16) and comparative 3D PCA plots grouping COLO 205 and MCF-7 datasets (figure 3.12 and supplementary figure for scree plots are S14 & S17). For the comparative 3D PCA PC1, PC2 and PC3 represented the maximum variance (95% and 99% in RAMAN and ATR-FTIR respectively) in both spectroscopic datasets. Both the analyses presented larger phase difference between TEI-PROSPR and UC-PROSPR, TEI and UC vectors being in close proximity to each other with small phase difference inferring about the correlation and variance in between groups. 3D PCA also presented that TEI and UC techniques for the two different sources had their nodal displacement in proximity with their datasets grouping in the same PC whereas PROSPR for each sample type had their nodes in different planes and its datasets group in different PC.

**Figure 3.12.**
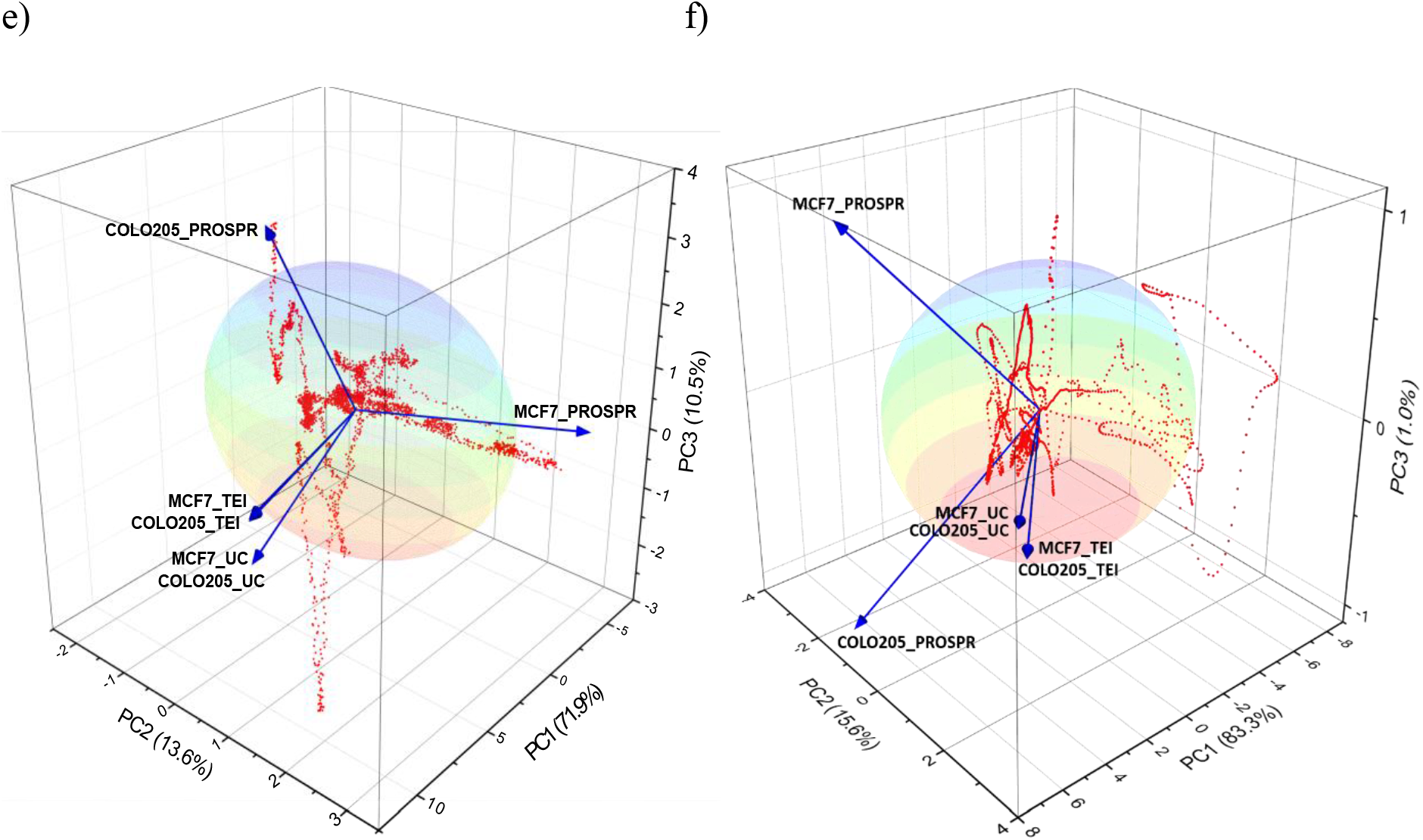
Multivariate analysis of RAMAN and ATR-FTIR spectra for differentiating the exosome isolation techniques and exosome types. PCA biplot for a) COLO 205 exosomes and b) MCF-7 exosomes analysis of RAMAN spectra differentiating between the techniques in the form of loading (in blue). The spectral data were represented as scores (in red). PCA biplot for c) COLO 205 exosomes and d) MCF-7 exosomes analysis of ATR-FTIR spectra differentiating between the techniques in the form of loading (in blue). The spectral data were represented as scores (in red). Comparative 3D PCA plots for e) RAMAN spectra and f) ATR-FTIR spectra of COLO 205 and MCF-7 exosomes differentiating between the techniques.

### Table of Quality Control (QC)

**Table of Quality Control (QC):**
| **Factors** | **TEI** | **PROSPR** | **UC** |
| --- | --- | --- | --- |
| **Yield of exosomes** | High | High | Intermediate |
| **Enrichment and post processing** | Enriched exosomes<br>No necessary post processing required | Low in enrichment<br>Post processing required to obtain pure exosomes | Enriched exosomes<br>Require refinement and purity with SEC or density gradient purity step. |
| **Morphology** | Good and evenly distributed | Average to undesirable | Good but presence of aggregation |
| **Protein profile** | High proteome recovery | Low protein recovery | Good recovery but low in comparison to TEI |
| **Zeta Potential and Stability** | High magnitude inferring better stability of particles | Intermediate magnitude. Have good stability | Low zeta potential values.<br>Incipient to moderate stability |
| **Determination of molecular components from RAMAN and FTIR spectral data** | Easy determination of spectra.<br>Proper separation in shifts of molecular components<br>Spectra with least standard deviation | Merged shifts or wavelengths that make it difficult to access and identify the components<br>Spectra with highest standard deviation | Easy and proper determination of molecular components<br>Spectral SD is in proximity to that of TEI |

## Conclusions

In our study, we compared the commonly used exosome isolation techniques TEI, PROSPR, and UC respectively. We analysed the exosomes isolated based on their morphological, biophysical and physiochemical characteristics by various methods to come to a conclusion about the usability of techniques based on the researcher’s application. Exosomes isolated by TEI had the highest yield compared to UC and PROSPR with no further requirement of post-processing for the enrichment of exosomes. PROSPR technique, on the other hand, had yielded exosomes and also some amount of microvesicles along with it that makes the technique require further post-processing for pure exosome enrichment. UC had enriched exosomes when compared to PROSPR but the presence of aggregating materials requires post-processing step a necessary one for this technique. Comparing these techniques based on the recovery and stability factors, TEI yielded good quality and well-dispersed exosomes with high stability compared to PROSPR which did not have expected morphology but had intermediate stability. UC had good recovery of morphology but the presence of aggregating particles resulted in low stability of exosomes in suspension making the particles agglomerate among themselves. PAGE analysis showed the TEI had better protein recovery compared to PROSPR and UC and the western blotting for the marker CD63 supported this observation. The determination of molecular composition through fingerprinting analysis was accurate and easy in the case of TEI followed by UC but was difficult in PROSPR spectra due to high randomness and merged spectral intensities. RAMAN and ATR-FTIR spectral analysis had similar trends for TEI and UC w.r.t statistical and multivariate analysis but differed in case of PROSPR. This may be due to the presence of non exosomal components like extracellular vesicles and apoptotic bodies contaminating the isolated exosomes. This finding was also supported with the determination of relative intensities of spectroscopic protein to lipid ratios.

In summary, TEI based isolation technique is the better technique to isolate exosomes that can be employed directly into clinical translation or other downstream applications. PROSPR can be advantageous to isolate EVs in general due to its low cost and ease of use but it could not be used for exosome with high purity and clinical applications. UC has been the conventional technique and is used frequently in research due to low cost of isolation but due to low particle stability and resultant smaller size because of high centrifugal force it’s not a suitable method for drug delivery or targeted therapeutics kind of applications.

## Acknowledgements

We thank the SRM Genetic Engineering facility, SRM Chemistry Research Facility for zeta potential, SRM Central Instrumentation Facility and technical staff for HRTEM and RAMAN spectroscopy, SRM Nanotechnology Research Centre and technical staff for ATR-FTIR, SRM Biotechnology Ultracentrifugation facility.

## Supplementary information

**Supplementary Figure S1.**
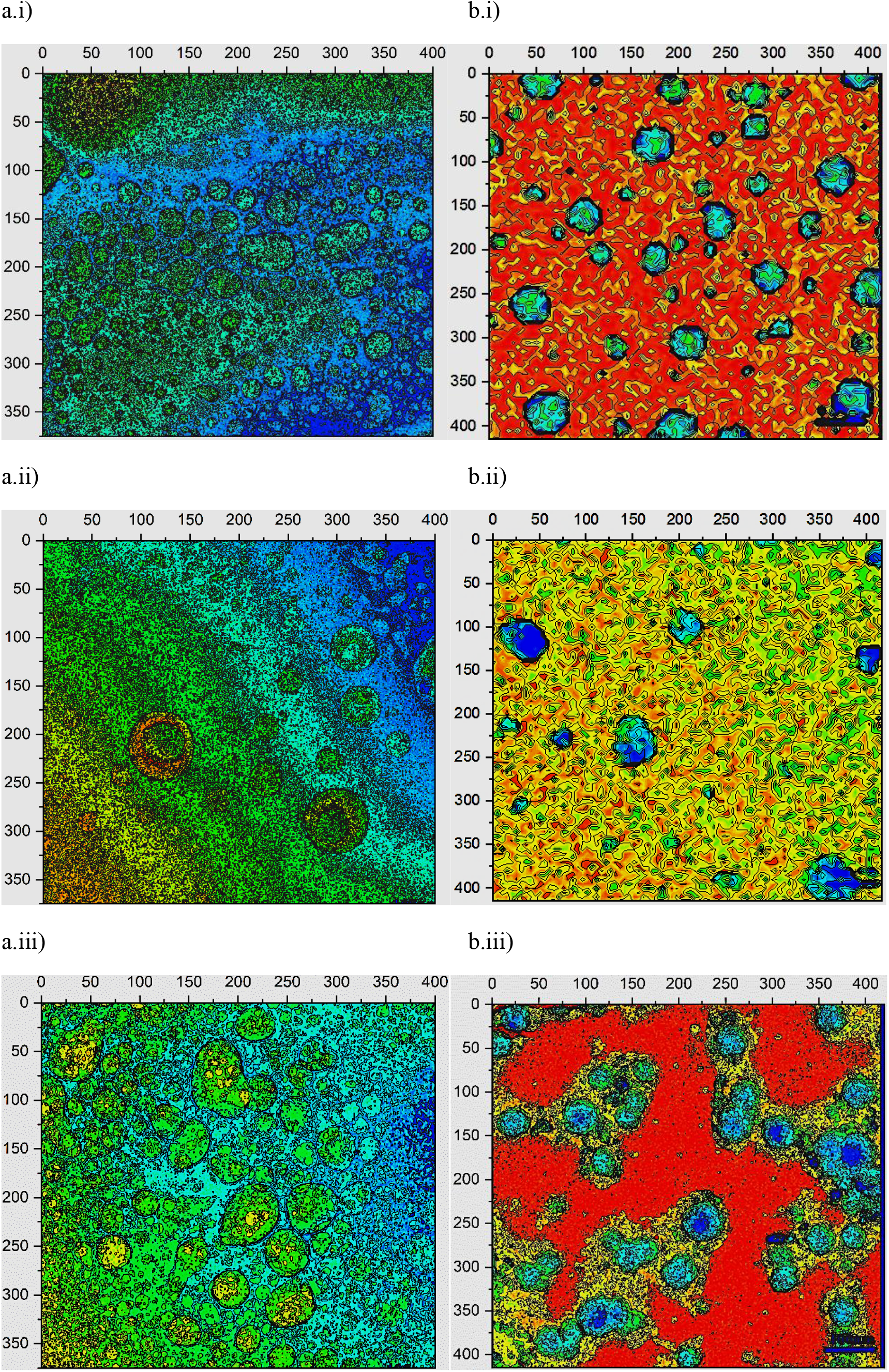
Representation of **contour maps plotted for the particle size distribution analysis of COLO 205 and MCF-7 exosome images obtained from HRTEM**. The analysis was done by calculating the exosome area, volume and size form contour line data that was extracted from the matrix of bitmap images created. a) COLO 205 exosomes isolated by i) TEI, ii) PROSPR and iii) UC. b) MCF-7 exosomes isolated by i) TEI, ii) PROSPR and iii) UC. The x and y axes were scaled with arbitrary units.

**Supplementary Figure S2.**
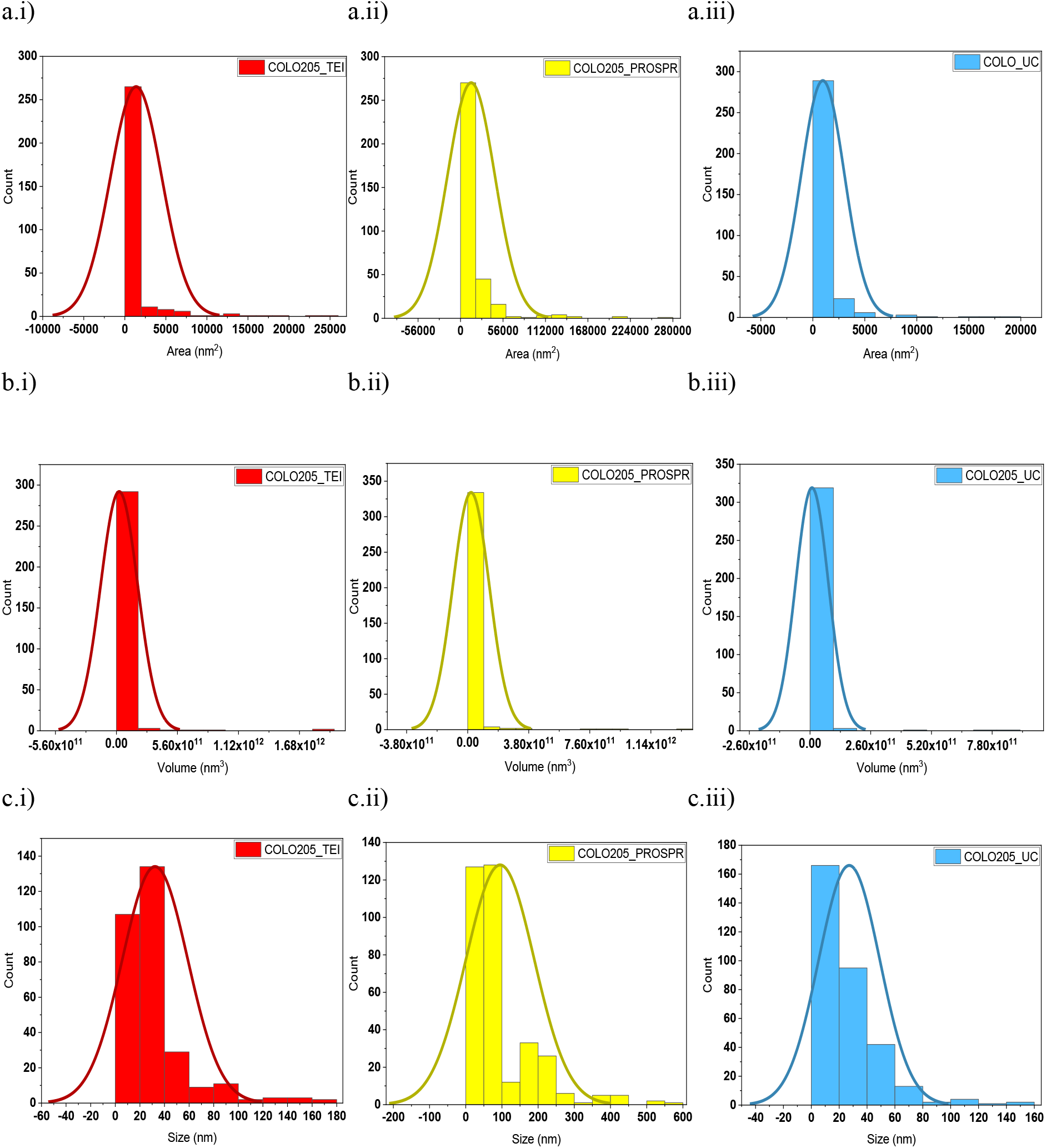
Individual normal distribution curves of COLO 205 exosomes. a) **Area** distribution curves of COLO 205 exosomes isolated by i) TEI, ii) PROSPR and iii) UC. b) **Volumetric** distribution curves of COLO 205 exosomes isolated by i) TEI, ii) PROSPR and iii) UC. c) **Size** distribution curves of COLO 205 exosomes isolated by i) TEI, ii) PROSPR and iii) UC.

**Supplementary Figure S3.**
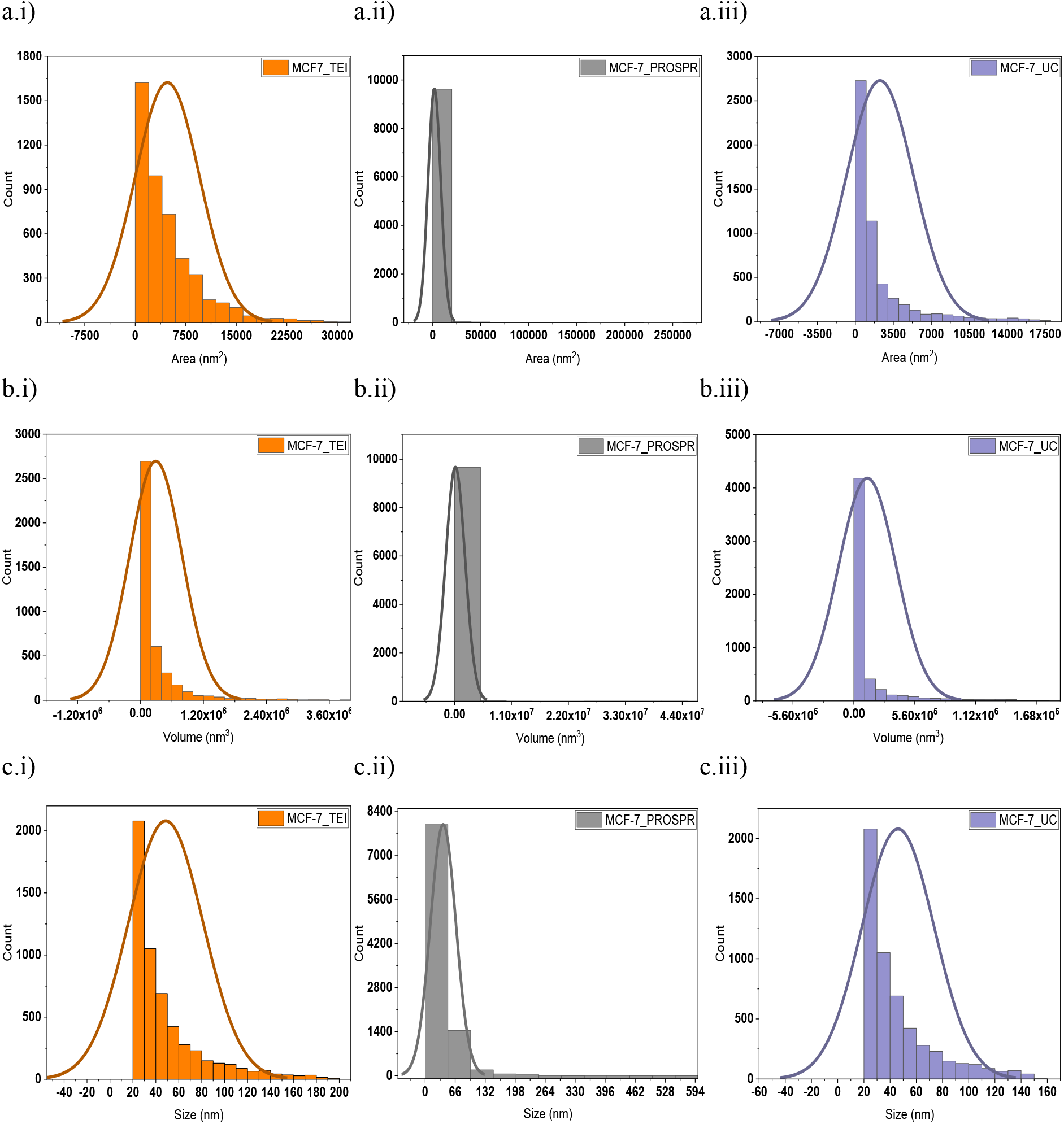
Individual normal distribution curves of MCF-7 exosomes. a) **Area** distribution curves of MCF-7 exosomes isolated by i) TEI, ii) PROSPR and iii) UC. b) **Volumetric** distribution curves of MCF-7 exosomes isolated by i) TEI, ii) PROSPR and iii) UC. c) **Size** distribution curves of MCF-7 exosomes isolated by i) TEI, ii) PROSPR and iii) UC.

**Supplementary Figure S4.**
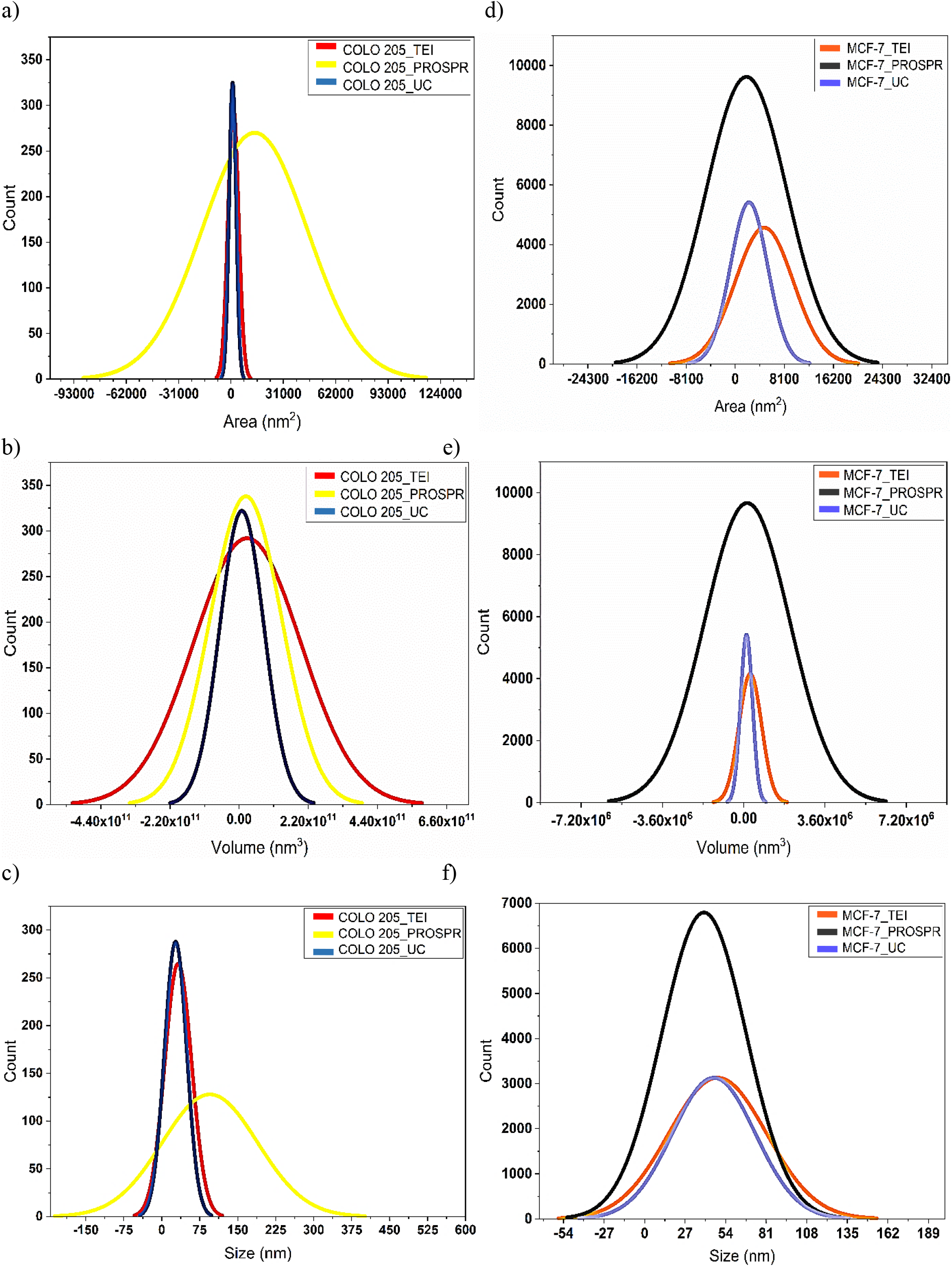
Comparative Normal Distribution curves of exosomes based on their area, volume and size as captured in HRTEM images. a-Area, b-Volumetric and c-Size distribution curves of COLO 205 exosomes. d-Area, e-Volumetric and f-Size distribution curves of MCF-7 exosomes.

**Supplementary Figure 5.**
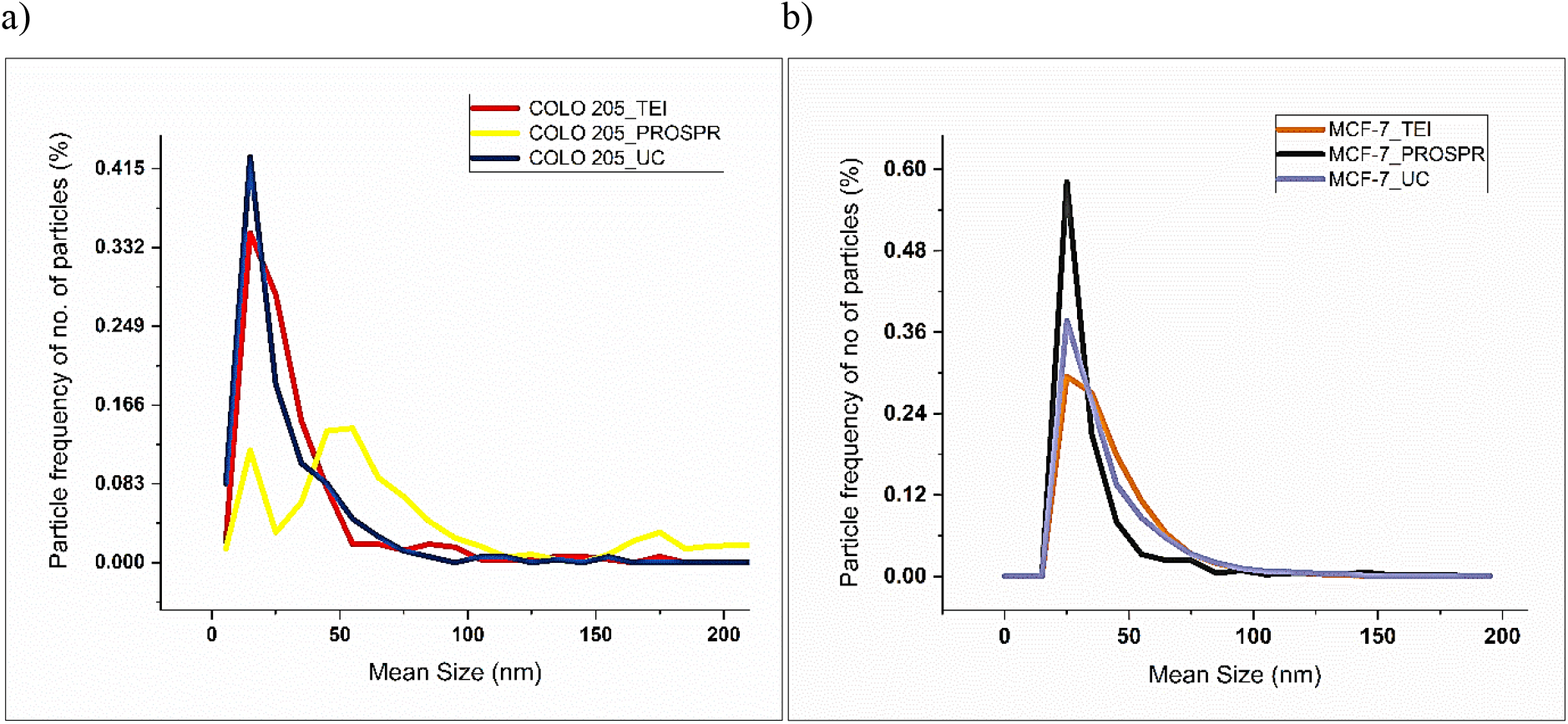
Frequency distribution curves of exosomes based on their size. a) Particle frequency curves of COLO 205 exosomes isolated by TEI, PROSPR, and UC for no. of particles based on their mean size. b) Particle frequency curves of MCF-7 exosomes isolated by TEI, PROSPR, and UC for no. of particles based on their mean size.

**Supplementary Figure 6.**
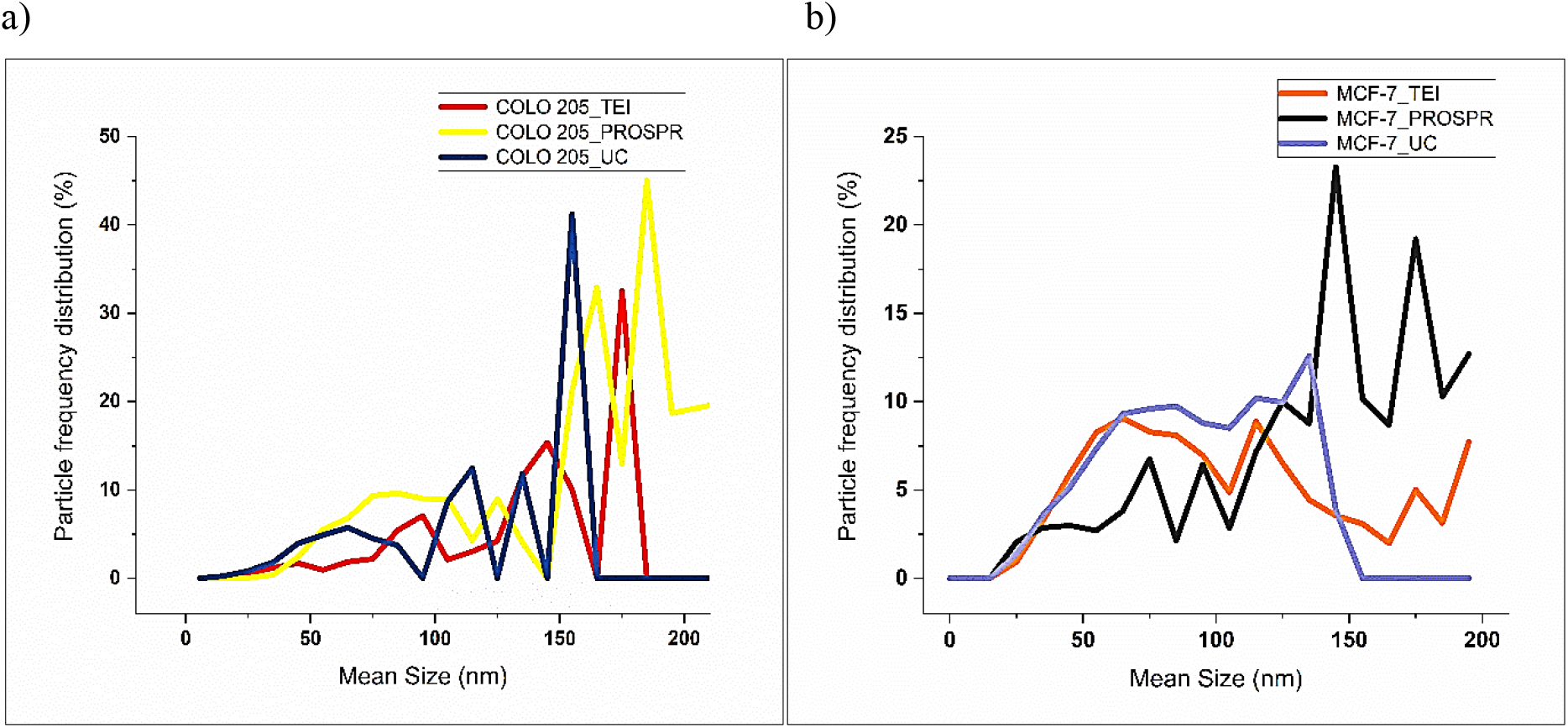
Frequency distribution curves of exosomes based on their size. a) Particle frequency distribution curves of COLO 205 exosomes isolated by TEI, PROSPR and UC for no. of particles based on their mean size. b) Particle frequency distribution curves of MCF-7 exosomes isolated by TEI, PROSPR and UC for no. of particles based on their mean size.

**Supplementary Table S1.** Quartile parameters of comparative box plots for COLO 205 exosomes.

| Parameters | TEI (nm) | PROSPR (nm) | UC (nm) |
| --- | --- | --- | --- |
| Min | 8.03 | 5.98 | 8.1 |
| First Quartile | 17.82 | 39.301 | 13.41 |
| Median Value | 24.14 | 54.42 | 20.73 |
| Third Quartile | 34.93 | 79.01 | 33.59 |
| Max | 174.82 | 667.2 | 154.51 |

**Supplementary Table S2.** Quartile parameters of comparative box plots for MCF-7 exosomes.

| Parameters | TEI (nm) | PROSPR (nm) | UC (nm) |
| --- | --- | --- | --- |
| Min | 22.56 | 22.56 | 20.28 |
| First Quartile | 37.42 | 25.23 | 26.92 |
| Median Value | 58.63 | 31.91 | 35.50 |
| Third Quartile | 84.44 | 43.70 | 54.31 |
| Max | 196.09 | 577.24 | 151.01 |

**Supplementary Figure S7.**
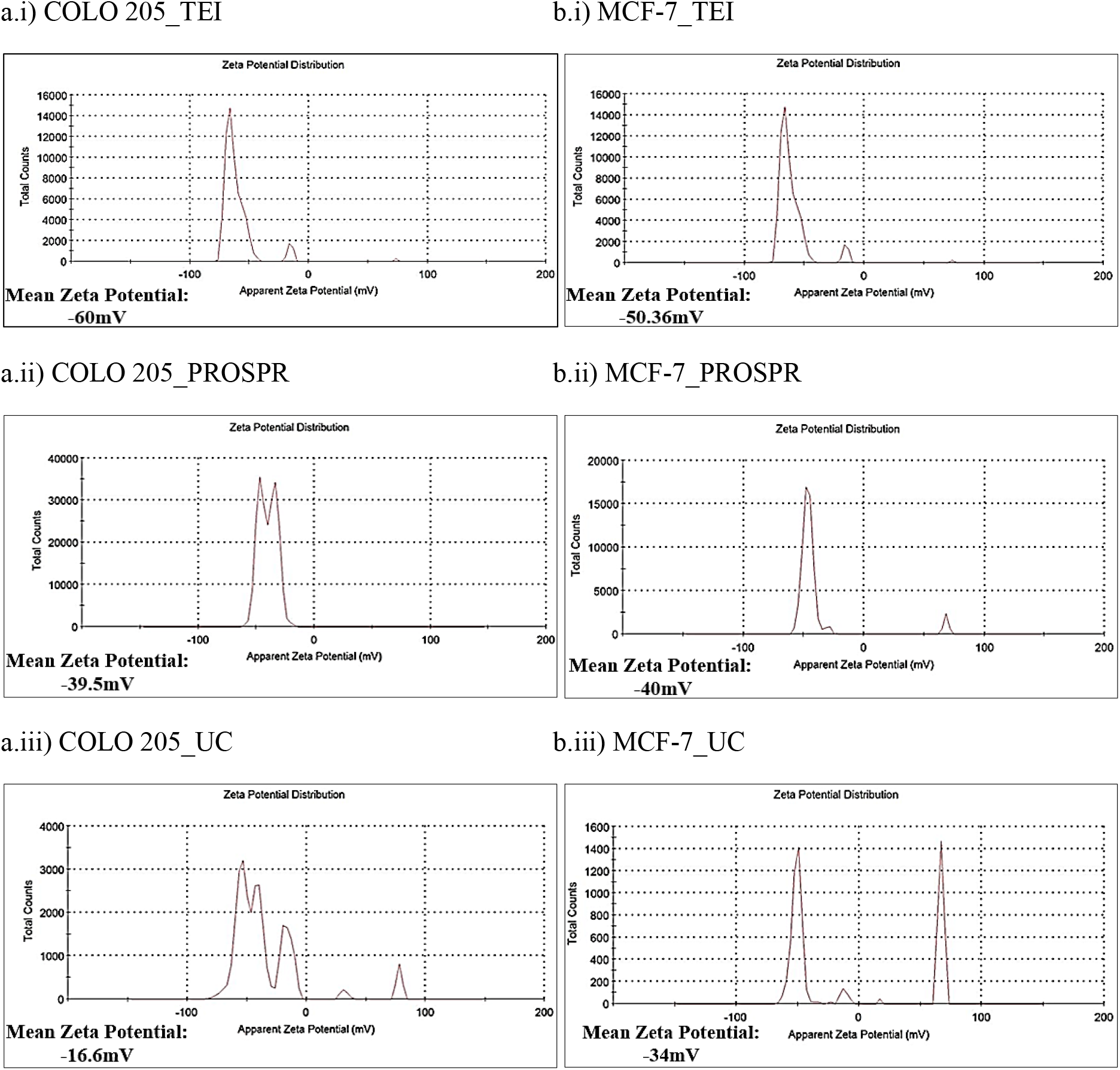
**System generated graphs for zeta potential analysis** of a) COLO 205 exosomes isolated by i) TEI ii) PROSPR and iii) UC. b) MCF-7 exosomes isolated by i) TEI ii) PROSPR and iii) UC. The representative graphs generated were average of three instrumental triplicates. The mean zeta potential was calculated by average of three biological triplicates.

**Supplementary Figure S8.**
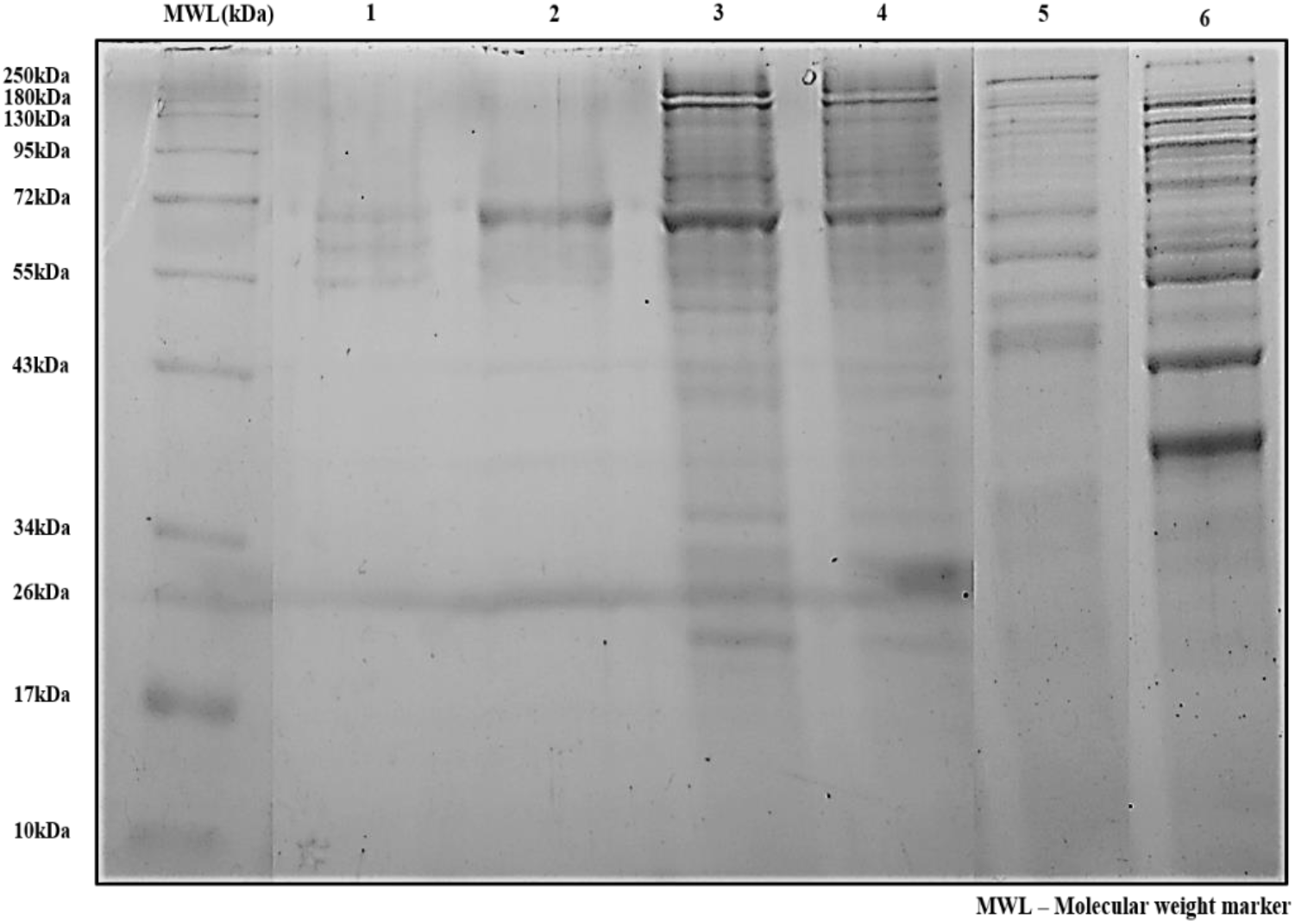
SDS-PAGE for comparative exosomal proteome profile. Sample loaded in lane 1 was exosomes isolated with TEI and treated with proteinase K. In lane 2 PROSPR sample, lane 3 COLO 205 exosomes isolate by TEI, lane 4 MCF-7 exosomes isolated by TEI, lane 5 UC samples and lane 6 total extracellular proteins isolated by acetone precipitation were loaded. The samples were directly digested and denatured with SDS sample loading buffer.

**Supplementary Figure S9.**
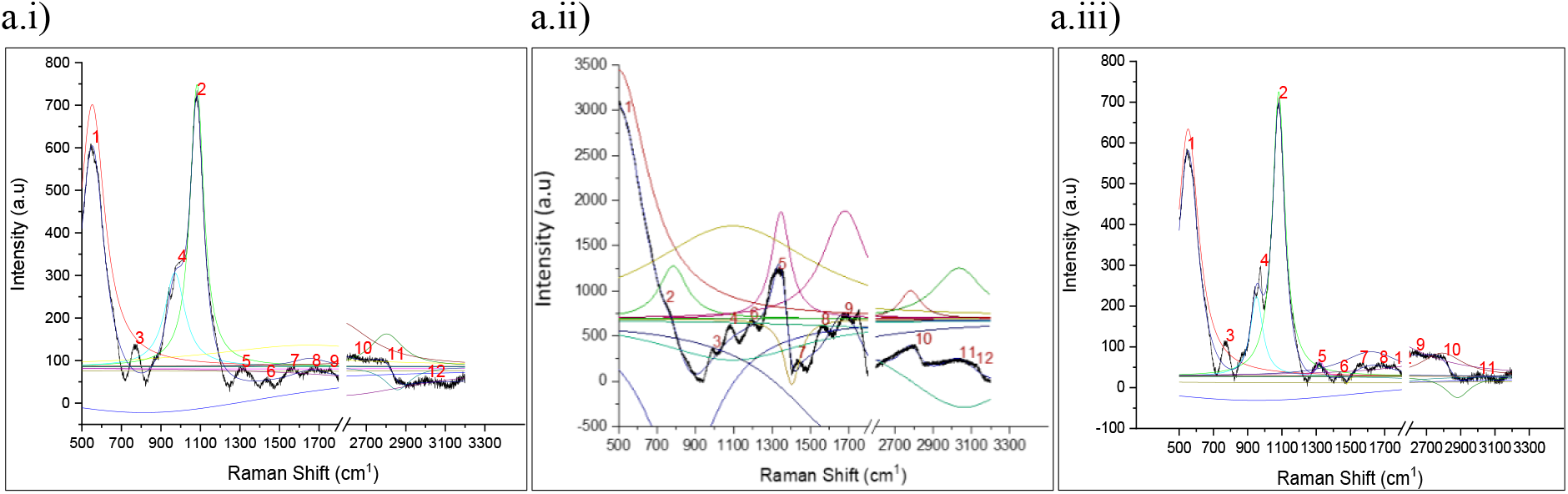
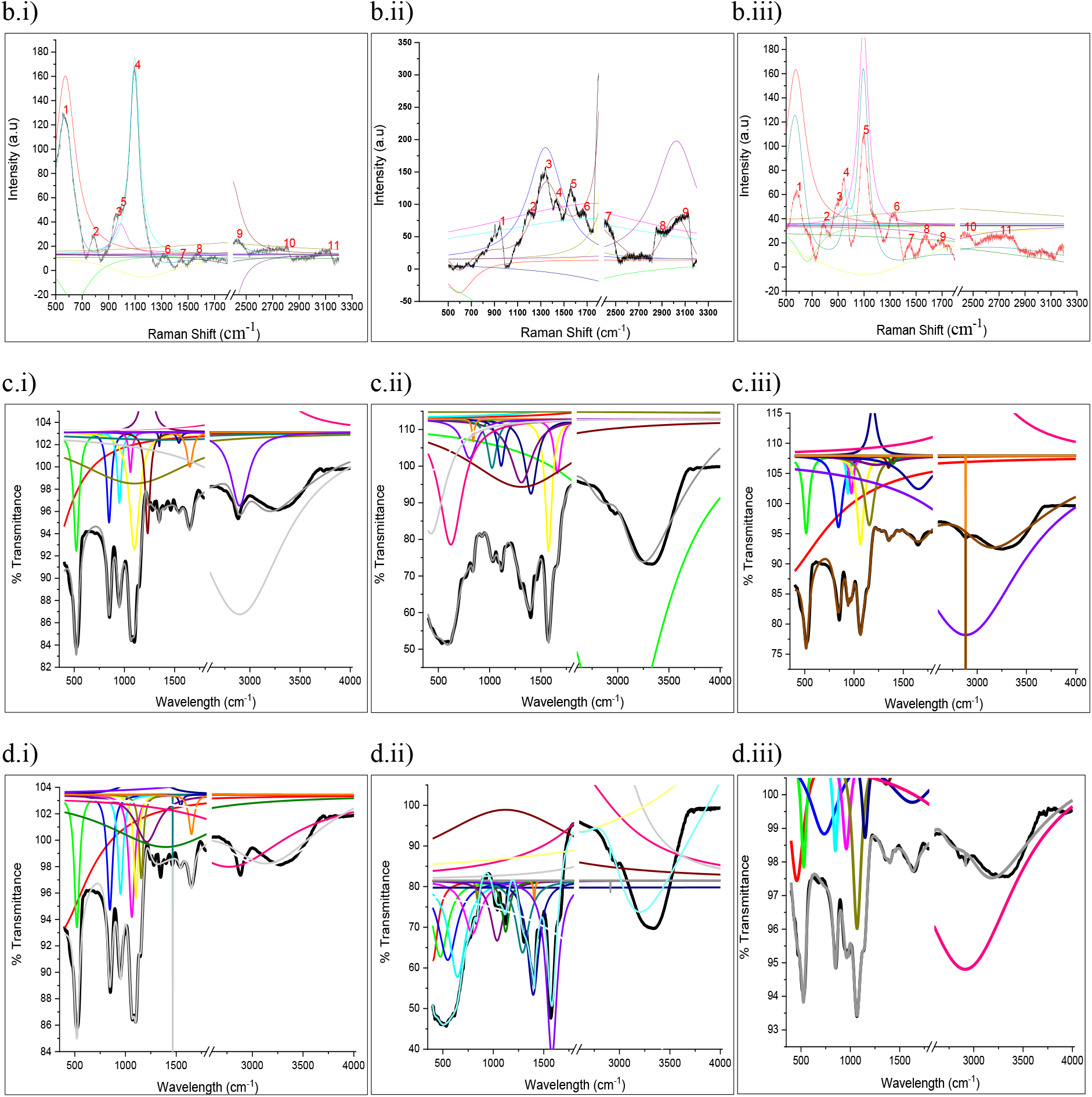
Representation of **non-linear multiple curve fitting and regression analyses performed for 400 iterations with Levenberg Marquardt algorithm (Function: Lorentz)** for determining the molecular components in **RAMAN spectra** of a) COLO 205 exosomes isolated by i) TEI with reduced chi sq. of 192 and R^2^ of 0.98, ii) PROSPR with reduced chi sq. of 739 and R^2^ of 0.97 and iii) UC with reduced chi sq. of 167 and R^2^ of 0.99. b) MCF-7 exosomes isolated by i) TEI with reduced chi sq. of 12 and R^2^ of 0.99, ii) PROSPR with reduced chi sq. of 39 and R^2^ of 0.97 and iii) UC with reduced chi sq. of 14 and R^2^ of 0.98. Non-linear multiple curve fitting and regression analyses were also performed for **ATR-FTIR spectra** for determining the molecular components. c) COLO 205 exosomes isolated by i) TEI with reduced chi sq. of 0.19 and R^2^ of 0.98, ii) PROSPR with reduced chi sq. of 2.68 and R^2^ of 0.98 and iii) UC with reduced chi sq. of 0.46 and R^2^ of 0.98. d) MCF-7 exosomes isolated by i) TEI with reduced chi sq. of 0.24 and R^2^ of 0.98, ii) PROSPR with reduced chi sq. of 7.68 and R^2^ of 0.97 and iii) UC with reduced chi sq. of 0.35 and R^2^ of 0.97. The RAMAN and ATR-FTIR spectra were deconvoluted for finding and retrieving the peaks withing mixture of spectrum containing the molecular components of exosomes.

**Supplementary Table S3.**
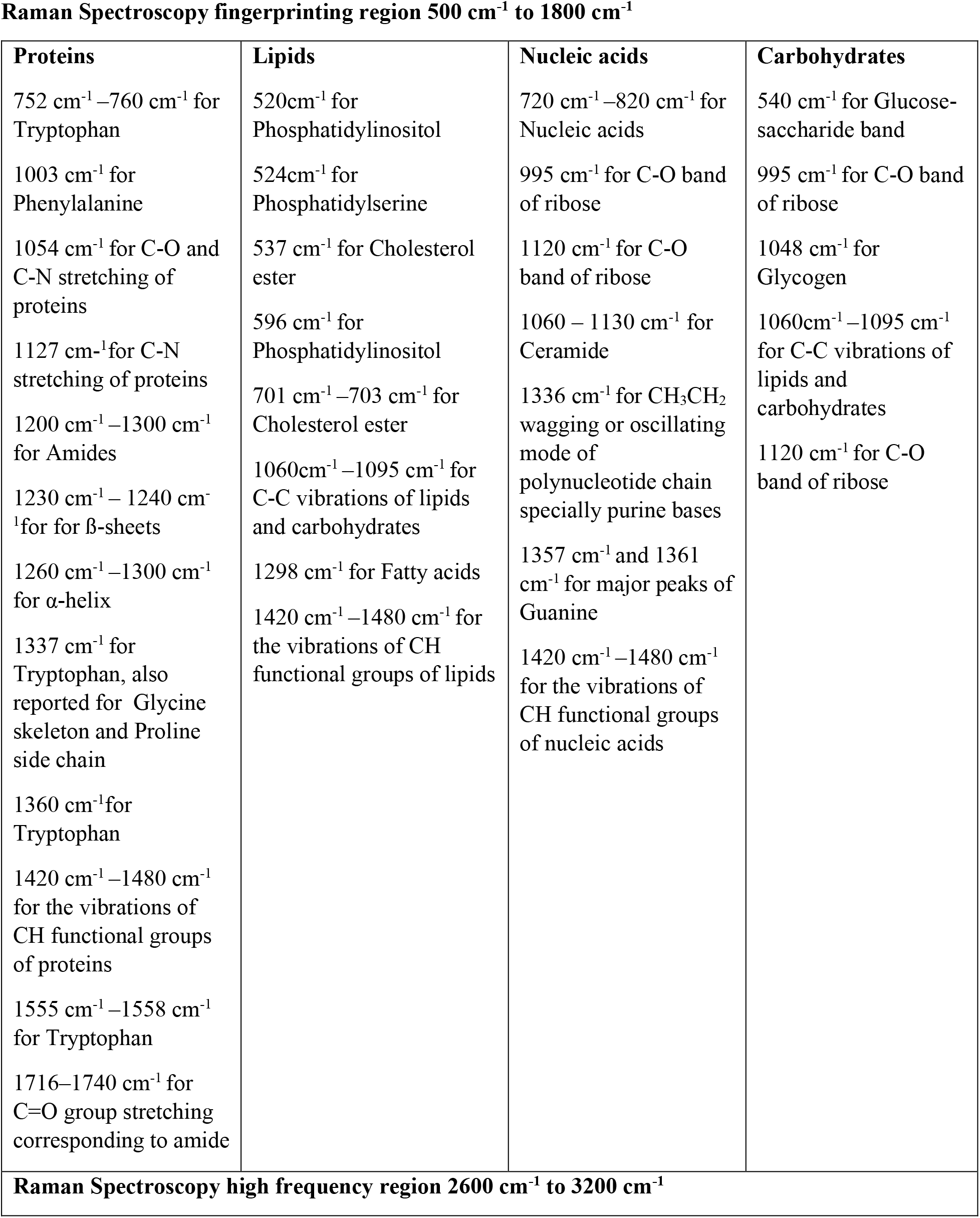
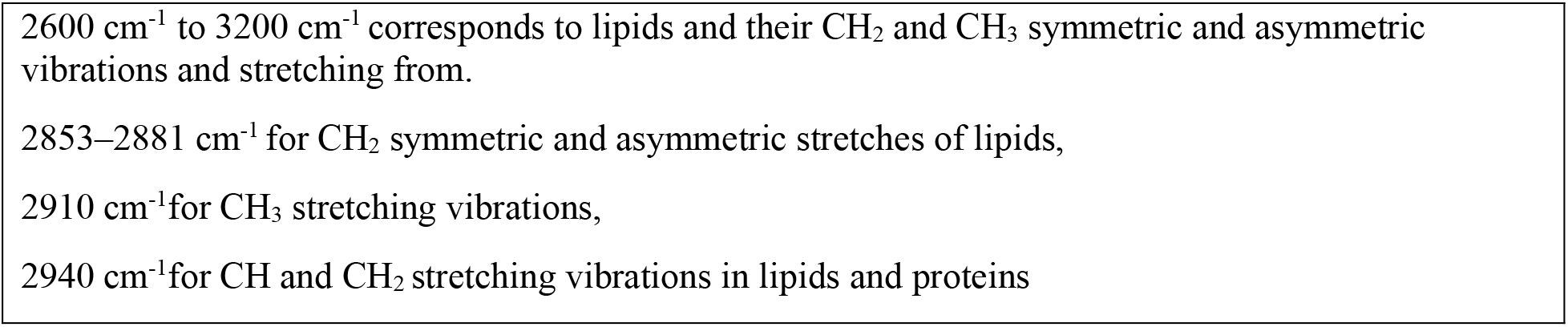
Exosomal components determined from RAMAN spectra of COLO 205 and MCF-7 after fitting of curves followed by deconvolution of spectra.

**Supplementary Table S4.**
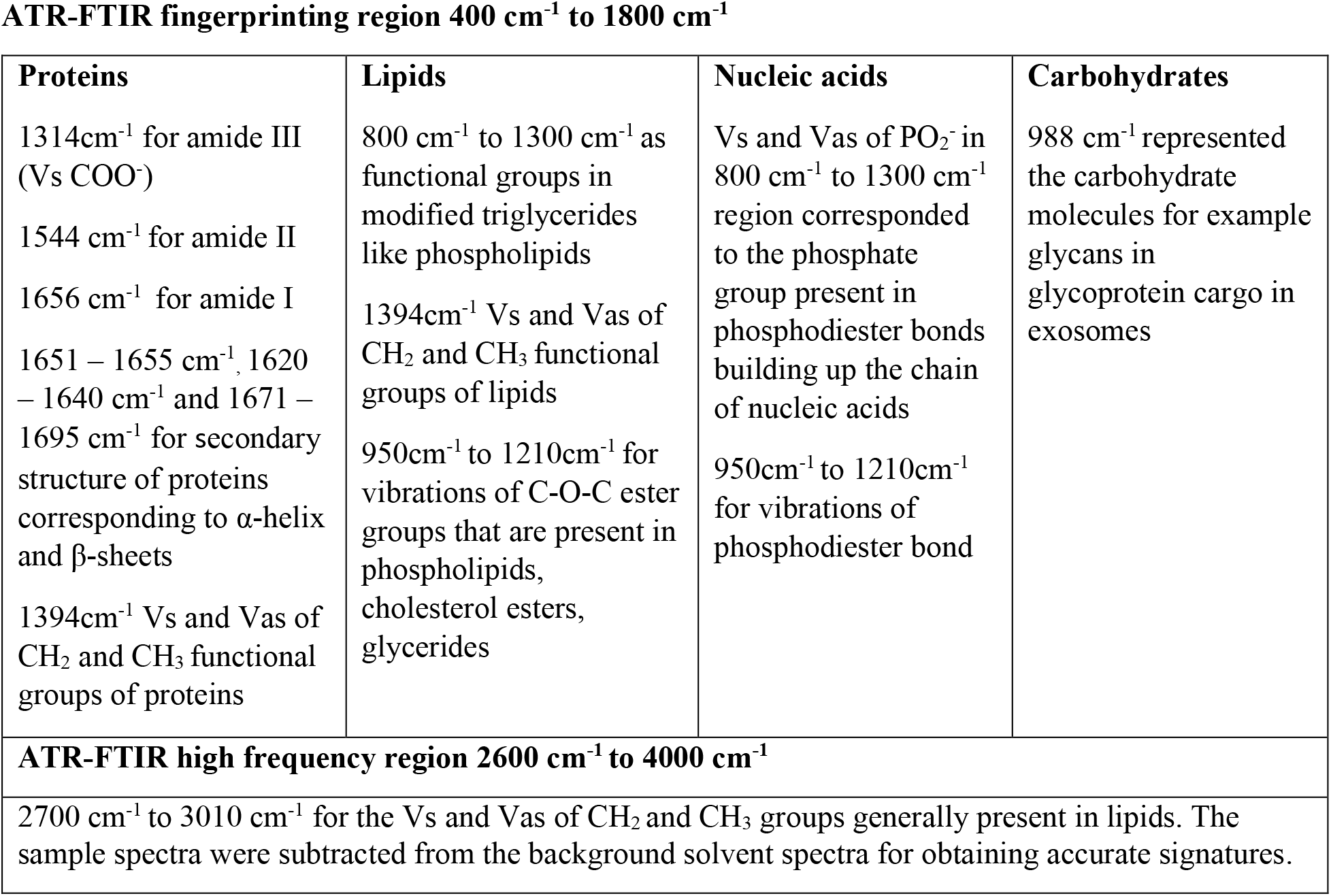
Exosomal components determined from ATR-FTIR spectra of COLO 205 and MCF-7 after fitting of curves followed by deconvolution of spectra.

**Supplementary Table S5.** Spectroscopic protein to lipid ratio calculated for a) **RAMAN spectra** and b) **ATR-FTIR spectra of COLO 205 and MCF-7 exosomes** isolated by TEI, PROSPR and UC.

| Isolation Techniques | Spectroscopic protein to lipid ratio calculated from ATR-FTIR spectra | | Mean ± SD |
| --- | --- | --- | --- |
|  | COLO 205 exosomes | MCF-7 exosomes |  |
| TEI | 1.184 | 1.184 | 1.18 ± 0.0001 |
| PROSPR | 1.484 | 1.51 | 1.49 ± 0.015 |
| UC | 1.18 | 1.176 | 1.18 ± 0.004 |

| Isolation Techniques | Spectroscopic protein to lipid ratio calculated from RAMAN spectra | | Mean ± SD |
| --- | --- | --- | --- |
|  | COLO 205 exosomes | MCF-7 exosomes |  |
| TEI | 1.142 | 0.985 | 1.06 ± 0.11 |
| PROSPR | 2.051 | 2.038 | 2.04 ± 0.01 |
| UC | 1.125 | 0.977 | 1.05 ± 0.1 |

**Supplementary Figure S10.**
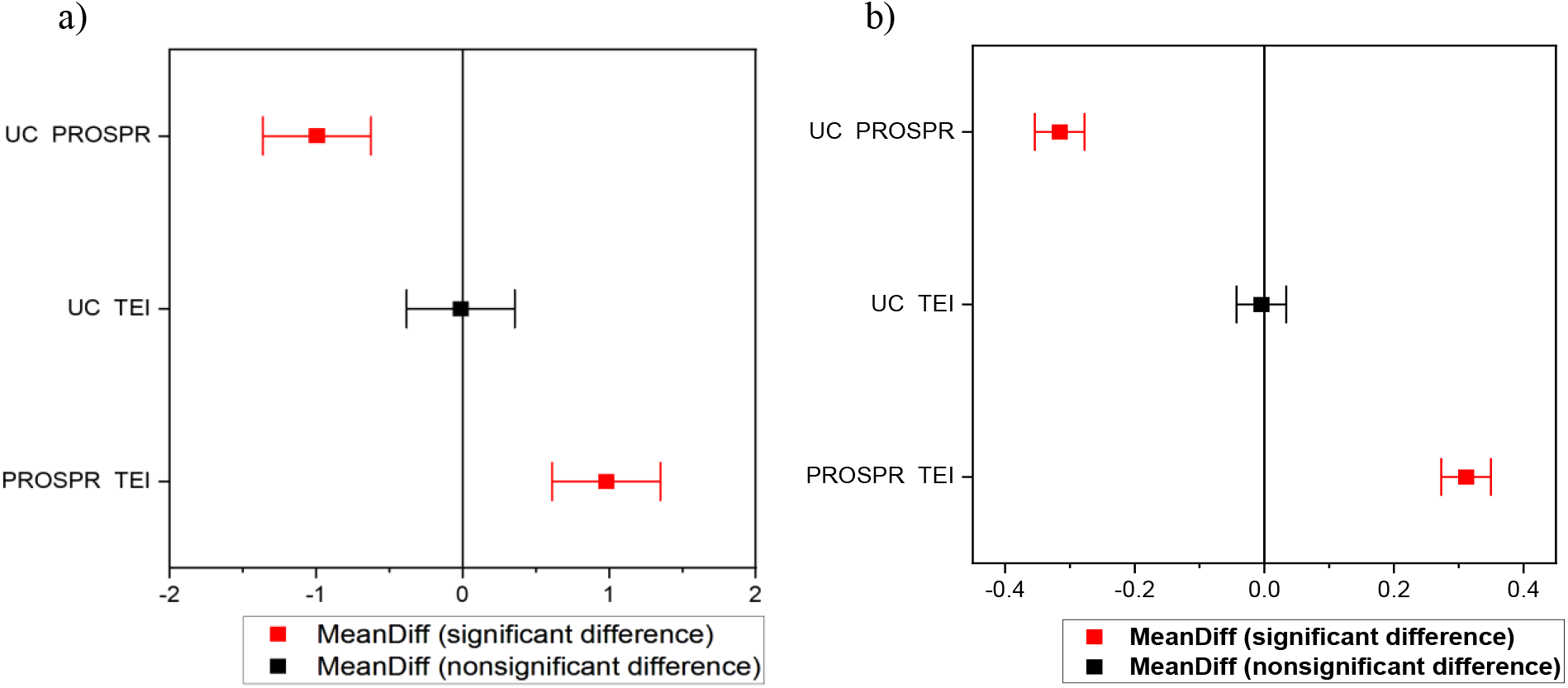
**Mean comparison plot** for Tukey’s test for the spectroscopic P/L ratio that was statistically analysed and validated with one-way ANOVA and by Tukey’s test for the relative intensities of a.ii) spectroscopic P/L ratio of RAMAN spectra and b.ii) spectroscopic P/L ratio of ATR-FTIR spectra.

**Supplementary Table S6.** a) Descriptive statistics b) Levene’s test (The Homogeneity of Variance tested at 0.05 level resulted in the population variances to be significantly different) c) One-way ANOVA (The null hypothesis assumed that the means of all levels were equal and the alternate hypothesis assumed that the means of one or more levels or groups were different. At 0.05 level, the population means were found to be significantly different) d) Tukey’s test (Significance equaled to 1 indicated that the difference of the means were significantly different at the 0.05 level but if the significance equaled to 0 that would have had indicated that the difference of the means were not significant at 0.05 level) **for RAMAN spectra of COLO 205 exosomes**. The statistical power was 1 for alpha 0.05 having sample size of 20022.

|  | N Analysis | N Missing | Mean | Standard Deviation | SE of Mean |
| --- | --- | --- | --- | --- | --- |
| COLO205_TEI | 6674 | 0 | 116.4 | 129.21 | 1.58 |
| COLO205_PROSPR | 6674 | 0 | 450.7 | 535.64 | 6.55 |
| COLO205_UC | 6674 | 0 | 91.48 | 127.5 | 1.56 |

b) Homogeneity of Variance Test (Levene's Test)
|  | DF | Sum of squares | Mean Square | F Value | Prob>F |
| --- | --- | --- | --- | --- | --- |
| Model | 2 | 301974000 | 150987000 | 2348.61 | 0 |
| Error | 20019 | 128698000 | 64287.73 |  |  |

c) One-way ANOVA (overall ANOVA)
|  | DF | Sum of squares | Mean Square | F Value | Prob>F |
| --- | --- | --- | --- | --- | --- |
| Model | 2 | 537056000 | 268528000 | 2518.55 | 0 |
| Error | 20019 | 2134420000 | 106619.9 |  |  |
| Total | 20021 | 2671480000 |  |  |  |

|  | MeanDiff | SEM | q Value | Prob | Alpha | Sig | LCL | UCL |
| --- | --- | --- | --- | --- | --- | --- | --- | --- |
| COLO205_PROSPR<br>COLO205_TEI | 334.3 | 5.65 | 89.9 | $3.3 * 10^{-16}$ | 0.05 | 1 | 345.97 | 372.46 |
| COLO205_UC<br>COLO205_TEI | -24.93 | 5.65 | 6.23 | $3.1 * 10^{-5}$ | 0.05 | 1 | 11.67 | 38.17 |
| COLO205_UC<br>COLO205_PROSPR | -359.21 | 5.65 | 83.63 | $3.3 * 10^{-16}$ | 0.05 | 1 | -348 | -321 |

a) Descriptive Statistics

**Supplementary Table S7.** a) Descriptive statistics b) Levene’s test (The Homogeneity of Variance tested at 0.05 level resulted in the population variances to be significantly different) c) One-way ANOVA (The null hypothesis assumed that the means of all levels were equal and the alternate hypothesis assumed that the means of one or more levels or groups were different. At 0.05 level, the population means were found to be significantly different) d) Tukey’s test (Significance equaled to 1 indicated that the difference of the means were significantly different at the 0.05 level but if the significance equaled to 0 that would have had indicated that the difference of the means were not significant at 0.05 level) for **RAMAN spectra of MCF-7 exosomes**. The statistical power was 1 for alpha 0.05 having sample size of 20022.

|  | N Analysis | N Missing | Mean | Standard Deviation | SE of Mean |
| --- | --- | --- | --- | --- | --- |
| MCF-7_TEI | 6674 | 0 | 29.93 | 28.1 | 0.34 |
| MCF-7_PROSPR | 6674 | 0 | 76.82 | 66.07 | 0.8 |
| MCF-7_UC | 6674 | 0 | 15.88 | 19.21 | 0.23 |

b) Homogeneity of Variance Test (Levene's Test)
|  | DF | Sum of squares | Mean Square | F Value | Prob>F |
| --- | --- | --- | --- | --- | --- |
| Model | 2 | 5327621.56 | 2663810.78 | 3076.32 | 0 |
| Error | 20019 | 17334600 | 865.9 |  |  |

c) One-way ANOVA (overall ANOVA)
|  | DF | Sum of squares | Mean Square | F Value | Prob>F |
| --- | --- | --- | --- | --- | --- |
| Model | 2 | 13591300 | 6795628.35 | 3690.24 | 0 |
| Error | 20019 | 36865200 | 1841.5 |  |  |
| Total | 20021 | 5.0456400 |  |  |  |

|  | MeanDiff | SEM | q Value | Prob | Alpha | Sig | LCL | UCL |
| --- | --- | --- | --- | --- | --- | --- | --- | --- |
| MCF-7_PROSPR<br>MCF-7_TEI | 46.89 | 0.74 | 89.26 | $3.3 * 10^{-16}$ | 0.05 | 1 | 45.14 | 48.62 |
| MCF-7_UC<br>MCF-7_TEI | -14.05 | 0.74 | 26.74 | $3.3 * 10^{-16}$ | 0.05 | 1 | -15.8 | -12.3 |
| MCF-7_UC<br>MCF-7_PROSPR | -60.93 | 0.74 | 116 | $3.3 * 10^{-16}$ | 0.05 | 1 | -62.6 | -59.19 |

a) Descriptive Statistics

**Supplementary Table S8.** a) Descriptive statistics b) Levene’s test (The Homogeneity of Variance tested at 0.05 level resulted in the population variances to be significantly different) c) One-way ANOVA (The null hypothesis assumed that the means of all levels were equal and the alternate hypothesis assumed that the means of one or more levels or groups were different. At 0.05 level, the population means were found to be significantly different) d) Tukey’s test (Significance equaled to 1 indicated that the difference of the means were significantly different at the 0.05 level and significance equaled to 0 indicated that the difference of the means were not significant at 0.05 level) for **ATR-FTIR spectra of COLO 205 exosomes**. The statistical power was 1 for alpha 0.05 having sample size of 20022.

|  | N Analysis | N Missing | Mean | Standard Deviation | SE of Mean |
| --- | --- | --- | --- | --- | --- |
| COLO205_TEI | 1868 | 0 | 98 | 3.53 | 0.08 |
| COLO205_PROSPR | 1868 | 0 | 83.15 | 14.41 | 0.33 |
| COLO205_UC | 1868 | 0 | 97.94 | 1.27 | 0.03 |

b) Homogeneity of Variance Test (Levene's Test)
|  | DF | Sum of squares | Mean Square | F Value | Prob>F |
| --- | --- | --- | --- | --- | --- |
| Model | 2 | 148015.43 | 74007.71 | 3892.16 | 0 |
| Error | 5601 | 106500.56 | 19.01 |  |  |

c) One-way ANOVA (overall ANOVA)
|  | DF | Sum of squares | Mean Square | F Value | Prob>F |
| --- | --- | --- | --- | --- | --- |
| Model | 2 | 273355.35 | 136677.67 | 1848.104 | 0 |
| Error | 5601 | 414225.26 | 73.95 |  |  |
| Total | 5603 | 687580.61 |  |  |  |

|  | MeanDiff | SEM | q Value | Prob | Alpha | Sig | LCL | UCL |
| --- | --- | --- | --- | --- | --- | --- | --- | --- |
| COLO205_PROSPR<br>COLO205_TEI | -14.84 | 0.28 | 74.6 | $3.3 * 10^{-16}$ | 0.05 | 1 | -15.5 | -14.18 |
| COLO205_UC<br>COLO205_TEI | -0.05 | 0.28 | 0.25 | 0.98 | 0.05 | 0 | -0.7 | 0.6 |
| COLO205_UC<br>COLO205_PROSPR | 14.8 | 0.28 | 74.33 | $3.3 * 10^{-16}$ | 0.05 | 1 | 14.13 | 15.45 |

a) Descriptive Statistics

**Supplementary Table S9.** a) Descriptive statistics b) Levene’s test (The Homogeneity of Variance tested at 0.05 level resulted in the population variances to be significantly different) c) One-way ANOVA (The null hypothesis assumed that the means of all levels were equal and the alternate hypothesis assumed that the means of one or more levels or groups were different. At 0.05 level, the population means were found to be significantly different) d) Tukey’s test (Significance equaled to 1 indicated that the difference of the means were significantly different at the 0.05 level but if the significance equaled to 0 that would have had indicated that the difference of the means were not significant at 0.05 level) for **ATR-FTIR spectra of MCF-7 exosomes**. The statistical power was 1 for alpha 0.05 having sample size of 20022.

|  | N Analysis | N Missing | Mean | Standard Deviation | SE of Mean |
| --- | --- | --- | --- | --- | --- |
| MCF-7_TEI | 1868 | 0 | 97.99 | 3.5 | 0.08 |
| MCF-7_PROSPR | 1868 | 0 | 82.01 | 15.72 | 0.36 |
| MCF-7_UC | 1868 | 0 | 97.94 | 1.27 | 0.03 |

b) Homogeneity of Variance Test (Levene's Test)
|  | DF | Sum of squares | Mean Square | F Value | Prob>F |
| --- | --- | --- | --- | --- | --- |
| Model | 2 | 171238.54 | 85619.27 | 3415.9 | 0 |
| Error | 5601 | 140388.46 | 25.06 |  |  |

c) One-way ANOVA (overall ANOVA)
|  | DF | Sum of squares | Mean Square | F Value | Prob>F |
| --- | --- | --- | --- | --- | --- |
| Model | 2 | 317082.46 | 158541.23 | 1818.41 | 0 |
| Error | 5601 | 488330.63 | 87.18 |  |  |
| Total | 5603 | 805413.09 |  |  |  |

|  | MeanDiff | SEM | q Value | Prob | Alpha | Sig | LCL | UCL |
| --- | --- | --- | --- | --- | --- | --- | --- | --- |
| MCF-7_PROSPR<br>MCF-7_TEI | -15.98 | 0.3 | 73.97 | $3.3 * 10^{-16}$ | 0.05 | 1 | -16.69 | -15.26 |
| MCF-7_UC<br>MCF-7_TEI | -0.04 | 0.3 | 0.23 | 0.98 | 0.05 | 0 | -0.7 | 0.66 |
| MCF-7_UC MCF-<br>7_PROSPR | 15.93 | 0.3 | 73.74 | $3.3 * 10^{-16}$ | 0.05 | 1 | 15.21 | 16.64 |

**Supplementary Figure 11.**
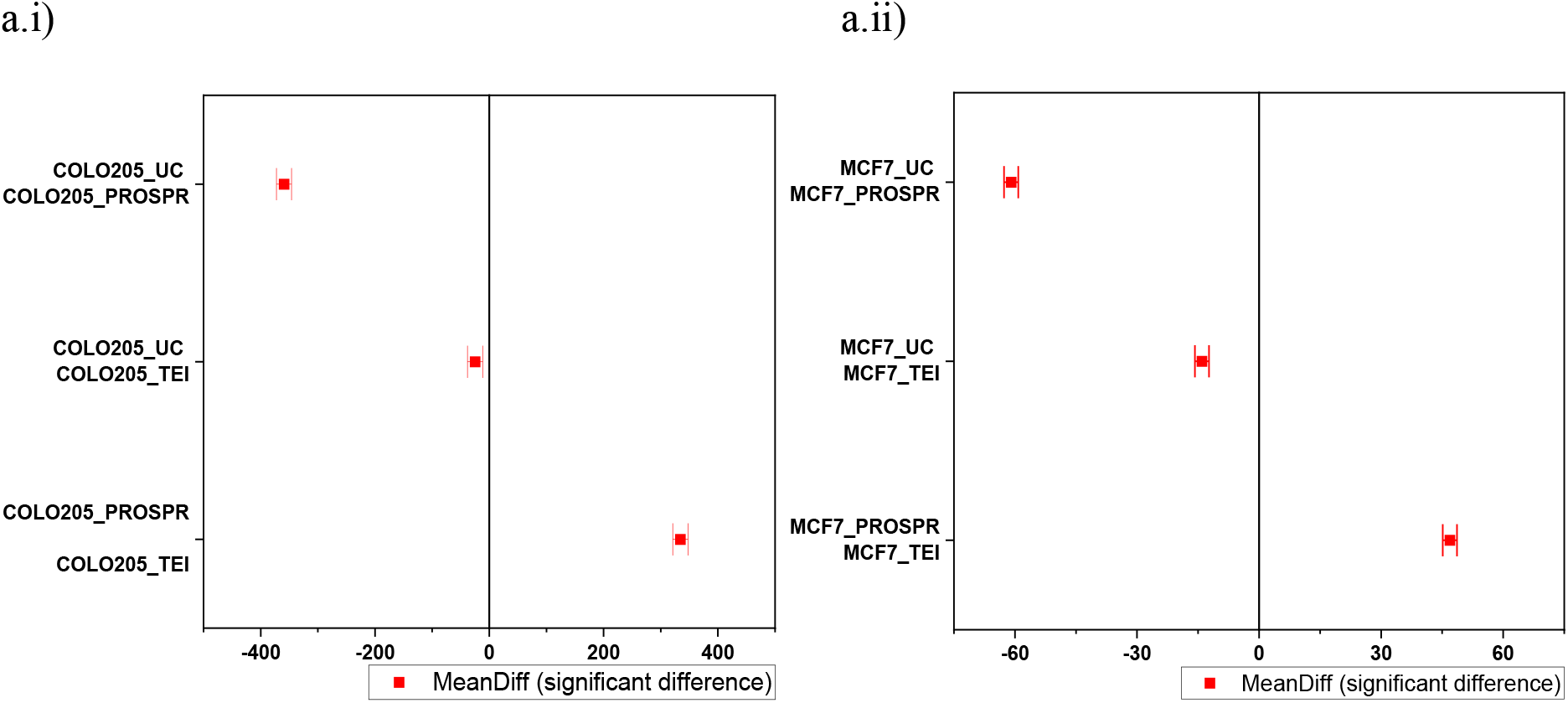
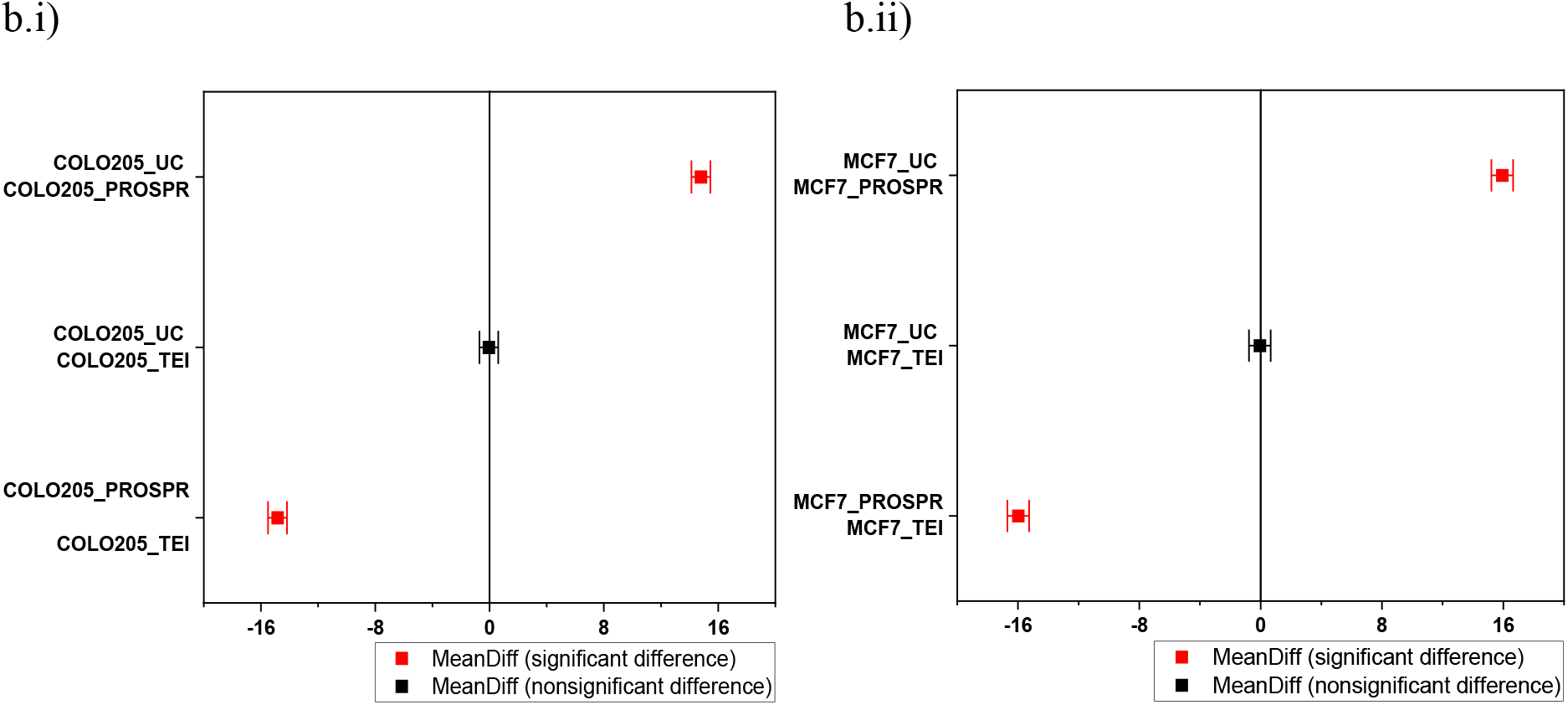
**Mean comparison plot for Tukey’s test** performed on the spectral data of a) RAMAN spectroscopy i) COLO 205 exosomes ii) MCF-7 exosomes and b) ATR-FTIR done for i) COLO 205 exosomes ii) MCF-7 exosomes. Mean comparison by Tukey’s test when had significance equals 1, it indicated that the difference of the means was significant at 0.05 level tested. If the significance equalled 0, it indicated that the difference of the means was not significant at 0.05 level tested.

**Supplementary Figure S12.**
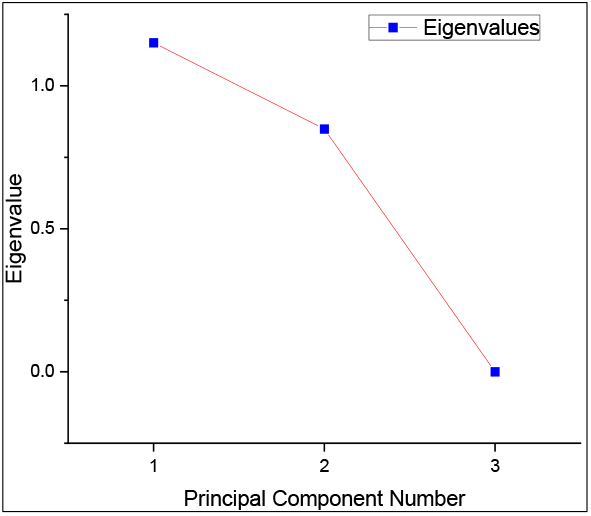
Scree plot for grouped PCA of RAMAN spectra for COLO 205 and MCF-7 exosomes.

**Supplementary Figure S13.**
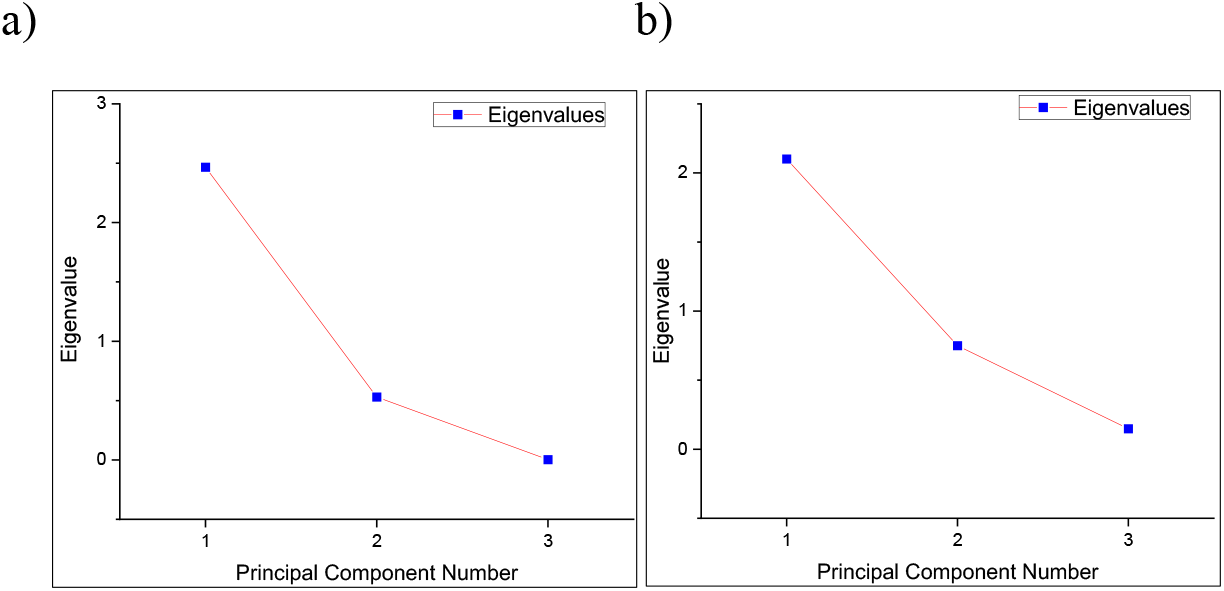
Scree plots for eigen values of PCA biplots plot for RAMAN spectra of a) COLO 205 exosomes and b) MCF-7 exosomes.

**Supplementary Figure S14.**
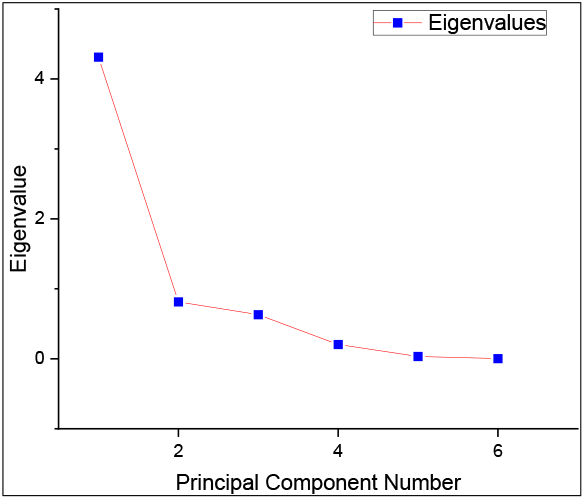
Scree plots for eigen values of 3D comparative PCA for RAMAN spectra of COLO 205 and MCF-7 exosomes.

**Supplementary Figure S15.**
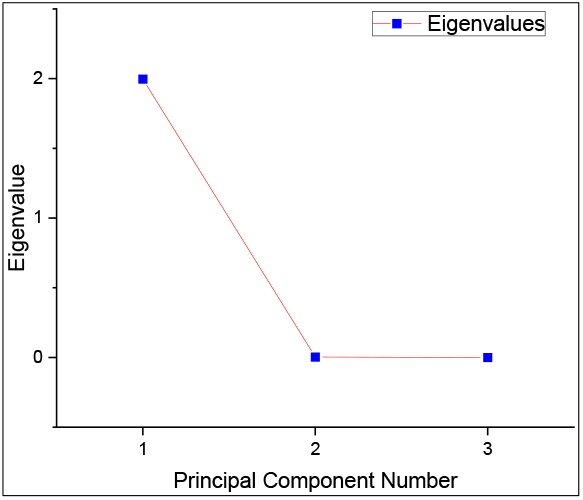
Scree plot for grouped PCA of ATR-FTIR spectra for COLO 205 and MCF-7 exosomes.

**Supplementary Figure S16.**
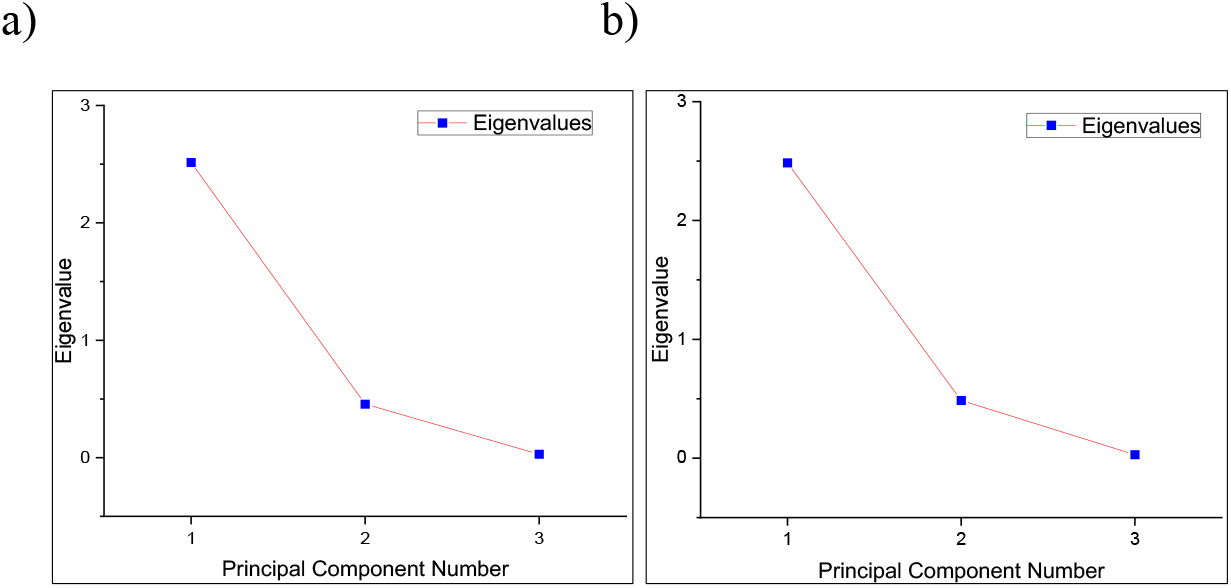
Scree plots for eigen values of PCA biplots plot for ATR-FTIR spectra of a) COLO 205 exosomes and b) MCF-7 exosomes.

**Supplementary Figure S17.**
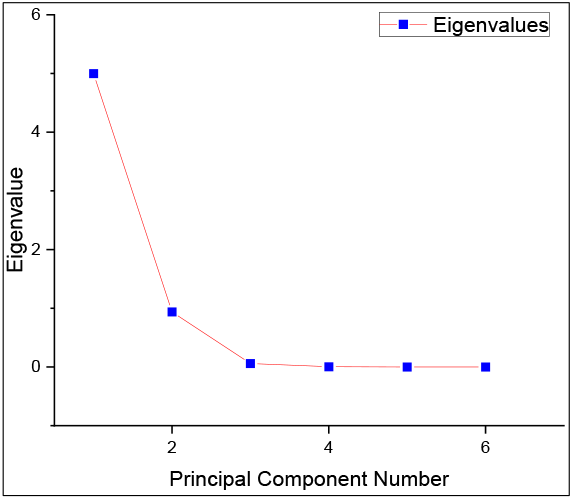
Scree plots for eigenvalues of 3D comparative PCA for ATR-FTIR spectra of COLO 205 and MCF-7 exosomes.

**Supplementary Figure 18.**
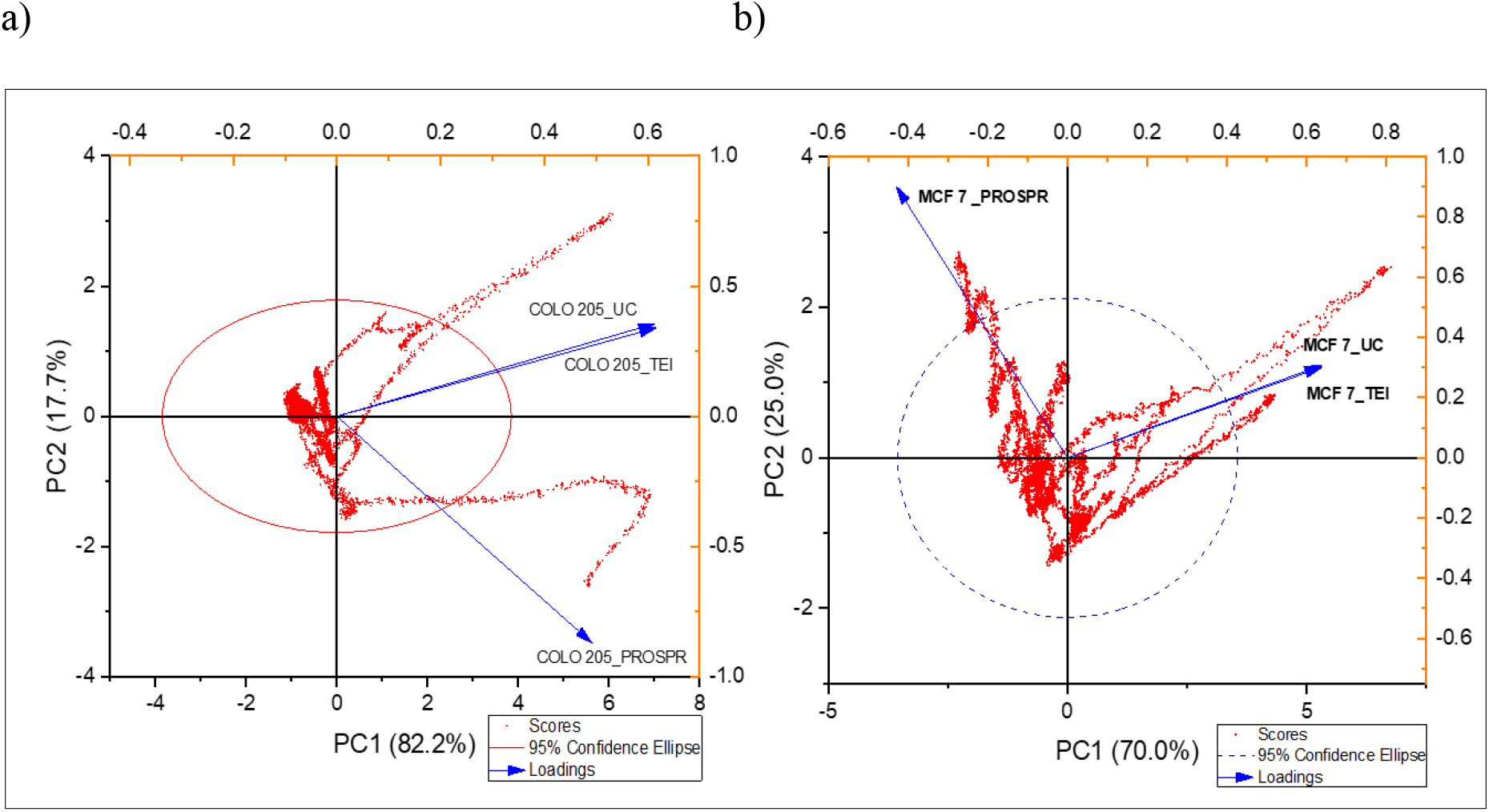
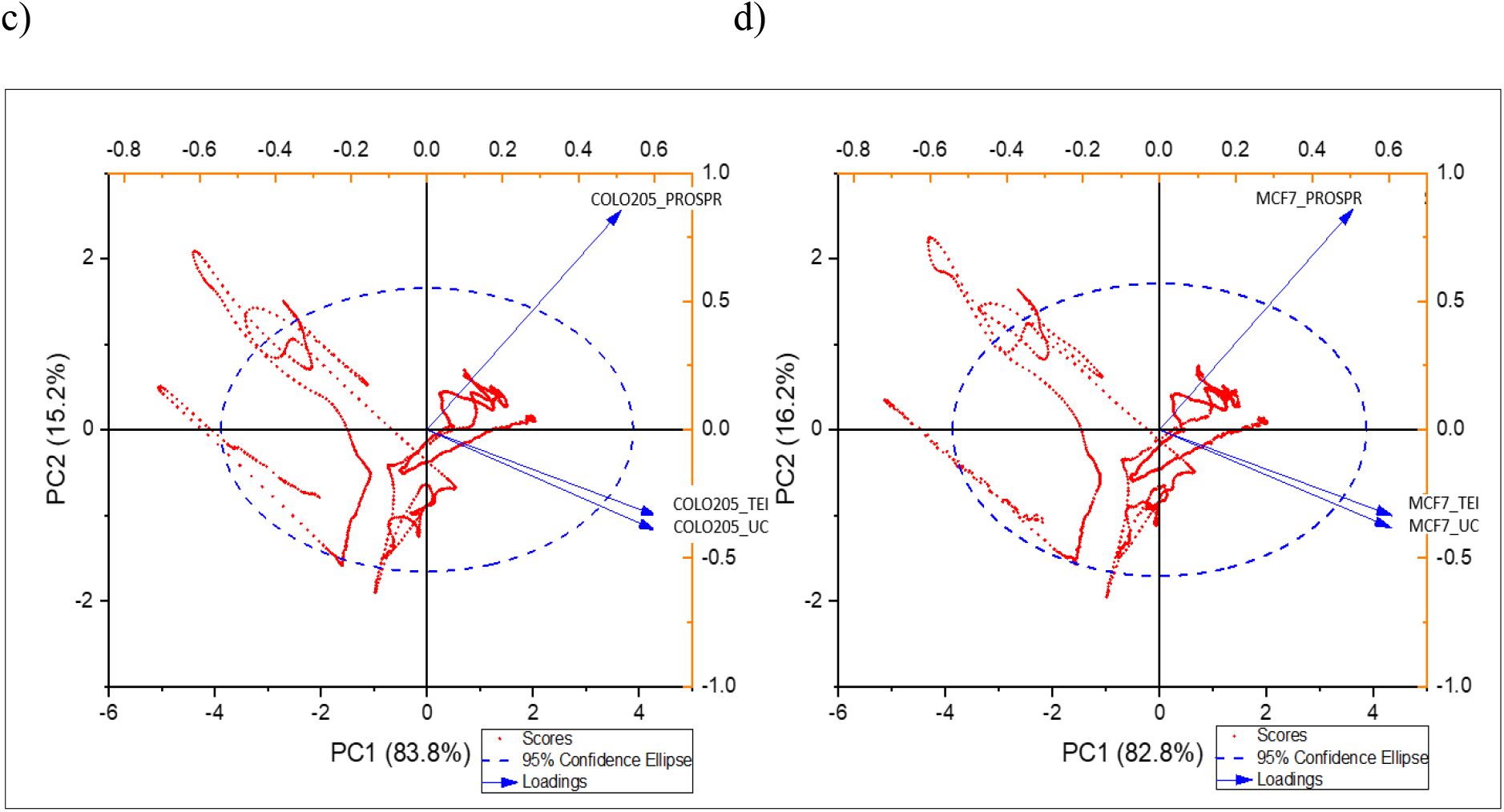
Multivariate analysis of RAMAN and ATR-FTIR spectra for differentiating the exosome isolation techniques and exosome types. **PCA biplot for a) COLO 205 exosomes and b) MCF-7 exosomes analysis** of RAMAN spectra differentiating between the techniques in the form of loading (in blue). The spectral data were represented as scores (in red). PCA biplot for c) COLO 205 exosomes and d) MCF-7 exosomes analysis of ATR-FTIR spectra differentiating between the techniques in the form of loading (in blue). The spectral data were represented as scores (in red). Comparative 3D PCA plots for e) RAMAN spectra and f) ATR-FTIR spectra of COLO 205 and MCF-7 exosomes differentiating between the techniques.

## References

1. Colombo, M., Raposo, G. & Théry, C. Biogenesis, Secretion, and Intercellular Interactions of Exosomes and Other Extracellular Vesicles. Annu. Rev. Cell Dev. Biol. (2014) doi:10.1146/annurev-cellbio-101512-122326.

2. Li, S. P., Lin, Z. X., Jiang, X. Y. & Yu, X. Y. Exosomal cargo-loading and synthetic exosome-mimics as potential therapeutic tools. Acta Pharmacologica Sinica (2018) doi:10.1038/aps.2017.178.

3. Zhang, L. & Yu, D. Exosomes in cancer development, metastasis, and immunity. Biochim. Biophys. Acta (BBA)-Reviews Cancer 1871, 455–468 (2019).

4. Willms, E. et al. Cells release subpopulations of exosomes with distinct molecular and biological properties. Sci. Rep. 6, 1–12 (2016).

5. Valadi, H. et al. Exosome-mediated transfer of mRNAs and microRNAs is a novel mechanism of genetic exchange between cells. Nat. Cell Biol. 9, 654–9 (2007).

6. Mathivanan, S., Ji, H. & Simpson, R. J. Exosomes: Extracellular organelles important in intercellular communication. J. Proteomics 73, 1907–1920 (2010).

7. Mashouri, L. et al. Exosomes: Composition, biogenesis, and mechanisms in cancer metastasis and drug resistance. Molecular Cancer (2019) doi:10.1186/s12943-019-0991-5.

8. Théry, C., Zitvogel, L. & Amigorena, S. Exosomes: Composition, biogenesis and function. Nature Reviews Immunology (2002) doi:10.1038/nri855.

9. Bunggulawa, E. J. et al. Recent advancements in the use of exosomes as drug delivery systems. J. Nanobiotechnology (2018) doi:10.1186/s12951-018-0403-9.

10. Zhang, M. et al. Exosome-based nanocarriers as bio-inspired and versatile vehicles for drug delivery: recent advances and challenges. Journal of Materials Chemistry B (2019) doi:10.1039/C9TB00170K.

11. Ha, D., Yang, N. & Nadithe, V. Exosomes as therapeutic drug carriers and delivery vehicles across biological membranes: current perspectives and future challenges. Acta Pharm. Sin. B 6, 287–296 (2016).

12. Balazs, D. A. & Godbey, W. Liposomes for Use in Gene Delivery. J. Drug Deliv. (2011) doi:10.1155/2011/326497.

13. Ma, B., Zhang, S., Jiang, H., Zhao, B. & Lv, H. Lipoplex morphologies and their influences on transfection efficiency in gene delivery. Journal of Controlled Release (2007) doi:10.1016/j.jconrel.2007.08.022.

14. Zhang, X. X., McIntosh, T. J. & Grinstaff, M. W. Functional lipids and lipoplexes for improved gene delivery. Biochimie (2012) doi:10.1016/j.biochi.2011.05.005.

15. Al-Dosari, M. S. & Gao, X. Nonviral gene delivery: Principle, limitations, and recent Progress. AAPS Journal (2009) doi:10.1208/s12248-009-9143-y.

16. Chen, W. C., May, J. P. & Li, S.-D. Immune responses of therapeutic lipid nanoparticles. Nanotechnol. Rev. 2, 201–213 (2013).

17. Liu, C. & Su, C. Design strategies and application progress of therapeutic exosomes. Theranostics (2019) doi:10.7150/thno.30853.

18. Antimisiaris, S. G., Mourtas, S. & Marazioti, A. Exosomes and exosome-inspired vesicles for targeted drug delivery. Pharmaceutics (2018) doi:10.3390/pharmaceutics10040218.

19. Lu, M. et al. Exosome-based small RNA delivery: Progress and prospects. Asian Journal of Pharmaceutical Sciences (2018) doi:10.1016/j.ajps.2017.07.008.

20. Li, X. et al. Challenges and opportunities in exosome research—Perspectives from biology, engineering, and cancer therapy. APL Bioeng. (2019) doi:10.1063/1.5087122.

21. Oves, M. et al. Exosomes: A paradigm in drug development against cancer and infectious diseases. J. Nanomater. 2018, (2018).

22. de la Torre Gomez, C., Goreham, R. V., Bech Serra, J. J., Nann, T. & Kussmann, M. ‘Exosomics’-A review of biophysics, biology and biochemistry of exosomes with a focus on human breast milk. Frontiers in Genetics (2018) doi:10.3389/fgene.2018.00092.

23. Colao, I. L., Corteling, R., Bracewell, D. & Wall, I. Manufacturing Exosomes: A Promising Therapeutic Platform. Trends in Molecular Medicine (2018) doi:10.1016/j.molmed.2018.01.006.

24. Zhang, Y., Shi, L. & Esfandiari, L. Biophysical Characterization of Exosomes Based on their Unique Dielectric Properties. Biophys. J. 118, 175a (2020).

25. Lobb, R. J. et al. Optimized exosome isolation protocol for cell culture supernatant and human plasma. J. Extracell. Vesicles 4, 1–11 (2015).

26. Rosenblum, D., Joshi, N., Tao, W., Karp, J. M. & Peer, D. Progress and challenges towards targeted delivery of cancer therapeutics. Nature Communications (2018) doi:10.1038/s41467-018-03705-y.

27. Patel, G. K. et al. Comparative analysis of exosome isolation methods using culture supernatant for optimum yield, purity and downstream applications. Sci. Rep. (2019) doi:10.1038/s41598-019-41800-2.

28. Yu, L. L. et al. A Comparison of Traditional and Novel Methods for the Separation of Exosomes from Human Samples. BioMed Research International (2018) doi:10.1155/2018/3634563.

29. Tang, Y. T. et al. Comparison of isolation methods of exosomes and exosomal RNA from cell culture medium and serum. Int. J. Mol. Med. (2017) doi:10.3892/ijmm.2017.3080.

30. Stranska, R. et al. Comparison of membrane affinity-based method with size-exclusion chromatography for isolation of exosome-like vesicles from human plasma. J. Transl. Med. (2018) doi:10.1186/s12967-017-1374-6.

31. Doyle, L. M. & Wang, M. Z. Overview of Extracellular Vesicles, Their Origin, Composition, Purpose, and Methods for Exosome Isolation and Analysis. Cells (2019) doi:10.3390/cells8070727.

32. Lobb, R. & Möller, A. Size exclusion chromatography: a simple and reliable method for exosome purification. in Extracellular Vesicles 105–110 (Springer, 2017).

33. Lee, J. et al. Enhanced paper-based ELISA for simultaneous EVs/exosome isolation and detection using streptavidin agarose-based immobilization. Analyst (2020) doi:10.1039/c9an01140d.

34. Fang, X. et al. Highly Efficient Exosome Isolation and Protein Analysis by an Integrated Nanomaterial-Based Platform. Anal. Chem. (2018) doi:10.1021/acs.analchem.7b04861.

35. Lim, J. et al. Direct isolation and characterization of circulating exosomes from biological samples using magnetic nanowires. J. Nanobiotechnology (2019) doi:10.1186/s12951-018-0433-3.

36. Clayton, A. et al. Analysis of antigen presenting cell derived exosomes, based on immuno-magnetic isolation and flow cytometry. J. Immunol. Methods (2001) doi:10.1016/S0022-1759(00)00321-5.

37. Oksvold, M. P., Neurauter, A. & Pedersen, K. W. Magnetic bead-based isolation of exosomes. Methods Mol. Biol. (2015) doi:10.1007/978-1-4939-1538-5_27.

38. Pedersen, K. W., Kierulf, B. & Neurauter, A. Specific and Generic Isolation of Extracellular Vesicles with Magnetic Beads. Methods Mol. Biol. (2017) doi:10.1007/978-1-4939-7253-1_7.

39. Helwa, I. et al. A comparative study of serum exosome isolation using differential ultracentrifugation and three commercial reagents. PLoS One 12, e0170628 (2017).

40. Contreras-Naranjo, J. C., Wu, H.-J. & Ugaz, V. M. Microfluidics for exosome isolation and analysis: enabling liquid biopsy for personalized medicine. Lab Chip 17, 3558–3577 (2017).

41. Liga, A., Vliegenthart, A. D. B., Oosthuyzen, W., Dear, J. W. & Kersaudy-Kerhoas, M. Exosome isolation: A microfluidic road-map. Lab Chip (2015) doi:10.1039/c5lc00240k.

42. Gallart-Palau, X. et al. Extracellular vesicles are rapidly purified from human plasma by PRotein Organic Solvent PRecipitation (PROSPR). Sci. Rep. (2015) doi:10.1038/srep14664.

43. Théry, C., Amigorena, S., Raposo, G. & Clayton, A. Isolation and Characterization of Exosomes from Cell Culture Supernatants and Biological Fluids. Curr. Protoc. Cell Biol. (2006) doi:10.1002/0471143030.cb0322s30.

44. Alcaraz, C., de Diego, M., Pastor, M. J. & Escribano, J. M. Comparison of a Radioimmunoprecipitation Assay to Immunoblotting and ELISA for Detection of Antibody to African Swine Fever Virus. J. Vet. Diagnostic Investig. (1990) doi:10.1177/104063879000200307.

45. Skliar, M. et al. Membrane proteins significantly restrict exosome mobility. Biochem. Biophys. Res. Commun. 501, 1055–1059 (2018).

46. Chernyshev, V. S. et al. Size and shape characterization of hydrated and desiccated exosomes. Anal. Bioanal. Chem. (2015) doi:10.1007/s00216-015-8535-3.

47. Jung, M. K. & Mun, J. Y. Sample Preparation and Imaging of Exosomes by Transmission Electron Microscopy. J. Vis. Exp. 5–9 (2018) doi:10.3791/56482.

48. Kotrbová, A. et al. TEM ExosomeAnalyzer: a computer-assisted software tool for quantitative evaluation of extracellular vesicles in transmission electron microscopy images. J. Extracell. Vesicles (2019) doi:10.1080/20013078.2018.1560808.

49. Linares, R., Tan, S., Gounou, C., Arraud, N. & Brisson, A. R. High-speed centrifugation induces aggregation of extracellular vesicles. Journal of Extracellular Vesicles (2015) doi:10.3402/jev.v4.29509.

50. Fernández-Llama, P. et al. Tamm-Horsfall protein and urinary exosome isolation. Kidney Int. (2010) doi:10.1038/ki.2009.550.

51. Wachalska, M. et al. Protein Complexes in Urine Interfere with Extracellular Vesicle Biomarker Studies. J. Circ. Biomarkers (2016) doi:10.5772/62579.

52. Gordiychuk, A., Svanera, M., Benini, S. & Poesio, P. Size distribution and Sauter mean diameter of micro bubbles for a Venturi type bubble generator. Experimental Thermal and Fluid Science (2016) doi:10.1016/j.expthermflusci.2015.08.014.

53. Yap, F. Y. et al. Quantitative morphometric analysis of hepatocellular carcinoma: Development of a programmed algorithm and preliminary application. Diagnostic Interv. Radiol. (2013) doi:10.4261/1305-3825.DIR.5973-12.1.

54. Kesimer, M. & Gupta, R. Physical characterization and profiling of airway epithelial derived exosomes using light scattering. Methods (2015) doi:10.1016/j.ymeth.2015.03.013.

55. Davis, C. N. et al. The importance of extracellular vesicle purification for downstream analysis: A comparison of differential centrifugation and size exclusion chromatography for helminth pathogens. PLoS Negl. Trop. Dis. (2019) doi:10.1371/journal.pntd.0007191.

56. Baker, M. J. et al. Using Fourier transform IR spectroscopy to analyze biological materials. Nat. Protoc. (2014) doi:10.1038/nprot.2014.110.

57. Smith, Z. J. et al. Single exosome study reveals subpopulations distributed among cell lines with variability related to membrane content. J. Extracell. Vesicles 4, 1–15 (2015).

58. Ali, S. M. et al. A comparison of Raman, FTIR and ATR-FTIR micro spectroscopy for imaging human skin tissue sections. Anal. Methods (2013) doi:10.1039/c3ay40185e.

59. Blat, A. et al. FTIR, Raman and AFM characterization of the clinically valid biochemical parameters of the thrombi in acute ischemic stroke. Sci. Rep. (2019) doi:10.1038/s41598-019-51932-0.

